# Mechanisms of vascular maturation and maintenance captured by longitudinal imaging of live mouse skin

**DOI:** 10.1101/2022.11.02.514907

**Authors:** Chen Yuan Kam, Ishani D. Singh, Catherine Matte-Martone, David G. Gonzalez, Paloma Solá, Guiomar Solanas, Júlia Bonjoch, Edward D. Marsh, Karen K. Hirschi, Valentina Greco

## Abstract

A functional network of blood vessels is essential for organ growth and homeostasis. Yet, how the vasculature matures and maintains adult homeostasis remains elusive in live mice. By longitudinally tracking the same neonatal endothelial cells (ECs) over days to weeks, we found that capillary plexus expansion is driven by network-wide vessel regression and transient angiogenesis. A fixed number of neonatal ECs rearrange their positions to evenly distribute throughout the developing plexus and become positionally stable in adulthood. Upon injury, while neonatal ECs are predisposed to die, adult ECs survive through a plasmalemmal self-repair response. Furthermore, adult neighboring ECs reactivate migration to assist vessel repair. Lastly, neonatal vessel regression and adult vascular maintenance are orchestrated by temporally restricted VEGFR2 dependent signaling. Our work sheds light on fundamental cellular mechanisms that underlie both vascular maturation and adult homeostasis *in vivo*.

## MAIN TEXT

Proper organ function requires an optimally constructed vascular network to ensure an adequate supply of nutrients and soluble factors via the bloodstream (*1*). This ability is particularly relevant to capillaries, the small diameter vessels (<10 microns) that function as the primary conduits for transport of soluble factors, nutrients, and immune cells to perfused tissues (*2*–*4*). Endothelial cells (ECs) are the functional units of capillaries, consisting of specialized squamous cells that form the inner lining of all blood vessels (*5*). ECs execute many processes involved in vascular network formation, refinement, and function by regulating fundamental aspects of vessel morphogenesis (*5*, *6*). Accordingly, much effort has been devoted to characterizing the behaviors of ECs in native *in vivo* environments. Investigations along these lines are crucial to advance our understanding of blood vessel structure and function in development, homeostasis, and disease.

The period immediately following birth is known as the neonatal developmental stage. It is an understudied, critical window that bridges the morphogenic events of embryogenesis to adult homeostasis (*7*–*9*). Our knowledge of *in vivo* vascular network formation has been greatly aided by the amenability of the embryonic zebrafish system to live imaging studies (*10*, *11*). Yet, there are significant gaps in our understanding of the tissue, cellular and molecular mechanisms involved in postnatal vessel network maturation and maintenance of adult vascular homeostasis. This is largely due to technical challenges in imaging zebrafish beyond embryonic stages, in addition to a loss of transparency in adulthood.

Skin, the largest organ in the human body, functions as an essential barrier to the external environment (*12*, *13*). Skin capillaries play a vital role in maintaining organ homeostasis: they control body temperature, regulate immune surveillance, and transport essential nutrients and paracrine factors (*14*–*16*). Here, we investigated cellular-, tissue-, and molecular-level mechanisms that define the maturation and maintenance of the mouse skin capillary plexus by adapting our previously established longitudinal intravital imaging approach (*17*). We found that regression and transient angiogenesis are central features of postnatal vessel maturation and that non-dividing ECs are vital players in the development of the skin’s vascular architecture. These cells grow and undergo positional changes necessary for tissue remodeling and regulation of local EC density. As the plexus reaches adulthood, remodeling decreases drastically, and ECs adopt a stable position. Adult, but not neonatal, ECs preferentially utilize a plasmalemmal self-repair process in response to injury, which maintains the stability of the skin vascular architecture. Collectively, our work demonstrates that distinct mechanisms regulate postnatal skin maturation and vascular homeostasis in adult skin, and that responses to injury also differ at the two developmental stages. Our findings may contribute to our understanding of pathologies and conditions characterized by vessel rarefaction or impaired vascular maintenance, such as scleroderma, primary hypertension, and aging (*18*–*20*). Furthermore, our results may form the basis for improved understanding of therapeutics that facilitate vessel maturation such as those that target the VEGF signaling pathway, and improve drug delivery during tumorigenesis (*21*).

### Vessel regression and transient angiogenesis drive maturation of the capillary network during skin plexus expansion

Capillaries are the primary conduits of the bloodstream to the tissues they perfuse (*2*–*4*). The mouse skin vasculature resides in the dermis and is organized in connected plexi throughout the organ (Supplementary Figure 1A & 1B). The most superficial capillary plexus is located directly below the epidermis (Supplementary Figure 1C). We sought to elucidate the mechanism by which the skin capillary plexus matures into an adult, architecturally defined network. To achieve this, we focused on the neonatal developmental period and visualized blood vessels in live mice using a new iteration of an intravital imaging technique established in our laboratory (*17*). We conducted our studies in palmoplantar skin as this anatomical region offers a simplified model devoid of cycling appendages (*22*, *23*).

We utilized the well-characterized *VE-CadherinCreER (VECadCreER*) mouse model to label the vascular architecture. This approach allowed for tamoxifen-inducible Cre-recombinase expression under control of the vascular endothelial cadherin (VECad) promoter (*24*). When crossed to Cre-dependent *mTmG* fluorescent reporter mice, tamoxifen administration induces a switch from membrane-tdTomato to membrane-EGFP expression, thus labeling all ECs with green fluorescence (Figure 1B) (*25*). Using this approach, we compared the skin superficial capillary plexus as it expands during various postnatal periods, including neonatal (P5), weanling (P21), and juvenile (P35) stages. We found that P5 mice exhibited significantly smaller capillary loops and more abundant branching points when compared to P21 cohorts (Figure 1A, 1A’ & 1A”). In addition, we observed an overall decrease in vascular coverage in the older mice (Figure 1A & 1A”’). When comparing P21 and P35 mice, these differences were not observed, indicating that the majority of postnatal vascular remodeling occurs before weaning (P21).

**Figure 1.**
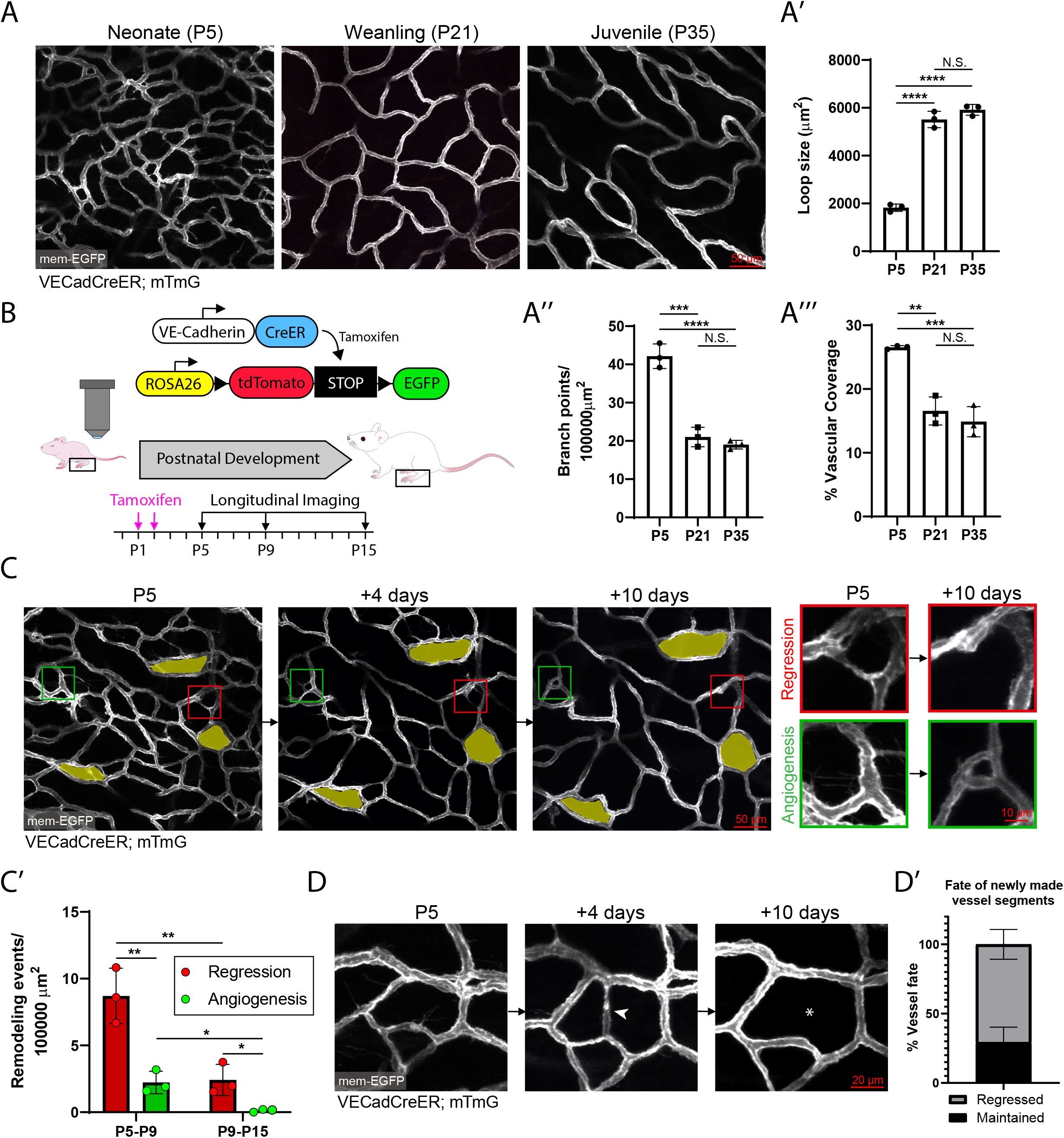
Vessel regression and transient angiogenesis drive maturation of the capillary network during postnatal skin plexus expansion. (A) Static comparison of the superficial capillary plexus of neonatal (P5), weanling (P21), and juvenile (P35) murine palmoplantar skin using fully induced *VECadCreER;mTmG* mice. (A’, A” & A”’) Quantification of capillary loop size (μm^2^), branching points, and % vascular coverage in P5, P21 and P35 animals (n=3 mice). N.S, not significant; **, P<0.01; ***, P<0.001; ****, P<0.0001 by one-way ANOVA followed by Tukey’s post-hoc test. (B) Schematic of the *VECadCreER;mTmG* tamoxifen-inducible fluorescent EC reporter and longitudinal imaging workflow. (C) Longitudinal imaging of the superficial capillary plexus of P5, P9 and P15 timepoints enables tracking of vessel remodeling events. The same capillary loops at successive time-points are pseudocolored in yellow. Regression events are defined as segments that are no longer detectable at later timepoints (Red inset) while angiogenesis events are new vessels formed at later time-points (Green inset). (C’) Quantification of overall vessel remodeling shows a significantly higher level of vessel regression compared to angiogenesis, both of which are further reduced in the P9-P15 interval compared to P5-P9 (n=3 mice). *, P<0.05; **, P<0.01 by unpaired Student’s t-test. (D) Tracking of vessels formed via angiogenesis (white arrowhead) at P5-P9 reveals that these new vessels are frequently pruned by the later time-point at P15. (D’) Quantification of the fate of angiogenesis events during P5-P9 and revisited at P15 shows that ~70% of new vessels go on to be regressed (n=38 events from 3 mice).

We next investigated the mechanisms responsible for the observed skin vascular changes during neonatal mouse development by establishing the ability to longitudinally track these same capillaries over time. We analyzed and revisited the superficial capillary plexus at P5, P9, and P15 and identified active remodeling events such as vessel regression, whereby initially present vessels were no longer detectable at subsequent time-points (Figure 1C; Red inset). In addition, we observed angiogenesis events in which new vessel segments sprouted from existing vessels (Figure 1C; Green inset). To assess the relative extent of regression versus angiogenesis during skin remodeling, we quantified these events over large areas (>300,000 μm^2^) during the P5-P15 developmental period. We found that vessel regression significantly outpaced angiogenesis at both the P5-P9 (4:1 ratio) and P9-P15 (18:1 ratio) intervals (Figure 1C’). Notably, overall vessel remodeling was more prevalent in P5-P9 compared to P9-P15 windows (~3-fold higher), indicating that the incidence of vessel remodeling decreases with postnatal time (Figure 1C’).

In accordance with previous studies (*26*), we find that vessels undergoing regression were indistinguishable from those undergoing angiogenesis as ~70% of these “sprouts” tracked from P5 to P9 resulted in regression compared to angiogenesis (Supplementary Figure 2A & 2A’). Additionally, P5 vessels exhibited widespread lateral filopodial extensions, established indicators of remodeling vessels, which significantly decrease as the network matures (Supplementary Figure 2B & 2B’) (*27*). To characterize the kinetics of vessel regression, we performed daily revisits and captured the spatiotemporal regulation of segment regression. We found that specific segments regressed within 1-2 days (Supplementary Figure 3A), while others followed slower kinetics, taking up to 4 days to complete the regression process (Supplementary Figure 3B). As such, we are able to demonstrate for the first time that the process of vessel regression displays relatively large kinetic variation.

**Figure 2.**
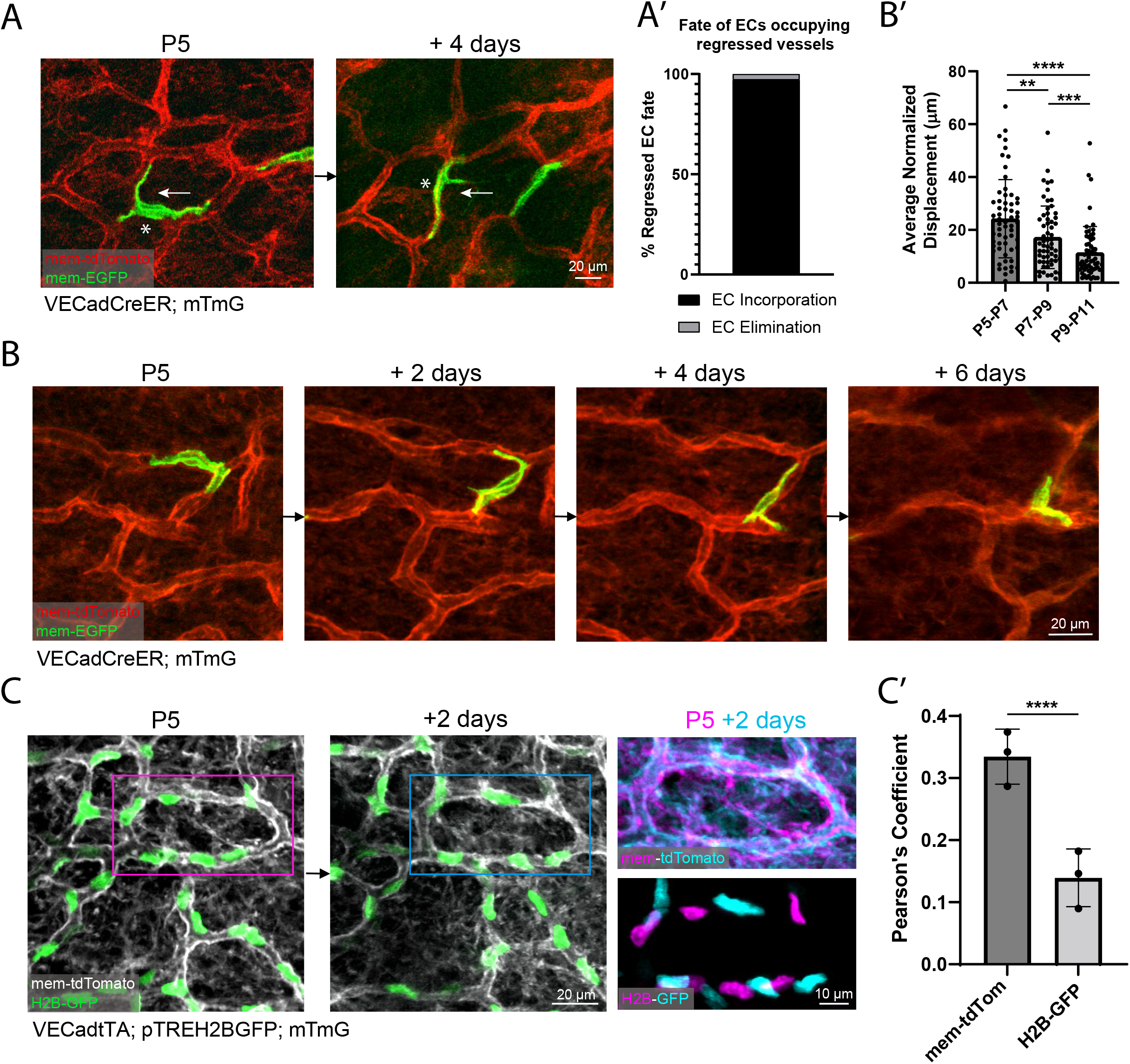
Neonatal ECs participate in plexus-wide positional rearrangement and execute selective vessel regression via a migratory mechanism. (A) Longitudinal tracking of single ECs in minimally induced *VECadCreER;mTmG* mice shows the execution of selective vessel regression via migration of the EC occupying the regressed segment (white asterisk). White arrow depicts direction of migration. (A’) Quantification of EC fate in vessels that are regressed shows the vast majority (37/38 events) of ECs are incorporated into adjacent vessels (n=38 cells from 5 mice). (B) Revisiting the same ECs within 2 day intervals reveals that ECs participate in migration within the existing vessel network architecture. (B’) Quantification of EC migration (average normalized displacement) calculated as changes in membrane occupancy between successive time-points indicates that the average rate of EC migration decreases over the course of postnatal development (n=60 cells from 3 mice). **, P<0.01; ***, P<0.001; ****, P<0.0001 by one-way ANOVA followed by Tukey’s post-hoc test. (C) Labeling of all EC nuclei with *VECadtTA;pTREH2BGFP* shows that EC rearrangement within the existing vascular architecture is a plexus-wide phenomenon. Overlay of EC positions from P5 (magenta) compared to 2 days later (cyan) indicates the extent of EC positional change. (C’) Quantification of the Pearson’s correlation coefficient between background corrected membrane signal (mem-tdTomato) compared to nuclear signal (H2BGFP) of P5 vs P7 vessels shows significantly higher correlation of membrane signal compared to nuclear signal (n=3 mice). ****, P<0.0001 by unpaired Student’s t-test.

Intriguingly, tracking of newly formed vessel segments (angiogenesis) during the P5-P9 interval showed that they were no longer detected when revisited at P15 (Figure 1D). Quantifications of the fates of all newly made branches revealed that approximately 70% of angiogenesis events are short-lived as they were found to be regressed at P15 (Figure 1D’). Collectively, our findings suggest that the cumulative vessel remodeling program of the dermal capillary plexus is driven by network-wide regression leading to a reduction of complexity and vascular coverage as postnatal development transitions into young adulthood.

### Neonatal ECs widely rearrange their positions within remodeling and stable capillary loops

We next sought to understand the cellular mechanisms underlying vessel regression. Our strategy to visualize single ECs and track their fates consisted of lowering the amount of tamoxifen (5 μg) administered to *VECadCreER; mTmG* mice (Figure 2A). We performed longitudinal imaging of single-labeled ECs over the postnatal period spanning P5-P15 and specifically monitored the fate of ECs in vessel segments that ultimately underwent regression. We successfully identified the same cells at successive time points in nearly all regression events (37/38); this observation indicates that EC apoptosis is not a significant contributor to vessel regression. Instead, we found that ECs migrated from regressing segments into non-remodeling vessels (Figure 2A & 2A’). Similar regression mechanisms have been described in zebrafish brain and mouse retinal vascular development (*26*, *28*, *29*),pointing to migration-mediated regression as a conserved developmental mechanism in vascular biology.

Most strikingly, we found that EC migration was broadly distributed and not restricted to regressing segments; indeed, we found evidence of EC migration in vessel loops that had not regressed (Figure 2B). As they changed positions, these ECs adapted their cellular morphology to the plexus structure, migrating within the endothelial layer. To assess the rate of EC migration, we developed a method to quantify changes in EC membrane occupancy based on the calculation of positional displacement at the leading and trailing edges of migrating ECs over time (Supplementary Figure 4). Analysis of EC migration over the postnatal P5-P11 interval revealed that the average rate of EC migration decreases over developmental time (Figure 2B’). We considered that our low induction method of labeling single ECs may be selective towards ECs expressing high levels of VE-Cadherin, and could therefore fail to capture an unbiased representation of the migratory ability of all ECs. Thus, we employed an independent strategy based on the use of a mouse reporter line (*VECadtTA;pTREH2BGFP;mTmG*) that allows for EC nuclei visualization and simultaneous tracking of all cells (Figure 2C) (*30*, *31*). Overlay of a conserved capillary loop at P5 and P7 revealed minimal positional overlap between P5 and P7 EC nuclei, confirming that the vast majority of capillary ECs undergo positional rearrangement during neonatal development (Figure 2C & 2C’). Together, our findings show that skin capillaries execute selective vessel regression through a migratory mechanism and that ECs undergo plexus-wide migration and rearrangement that decrease throughout neonatal development.

### Cellular density in the developing plexus is locally controlled by the incorporation of ECs from regressed segments

The widespread movement of ECs during skin development suggests that local cell density might be affected as cells from regressing segments are incorporated into nearby capillary loops. To test this, we used longitudinal tracking of P5 and P21 mice to monitor cellular density as the remodeling plexus expanded and reached maturity. We hypothesized that capillary loops containing regressed segments would exhibit a larger increase in cell number per area unit than conserved loops without regressions. We determined the net change in EC number by longitudinally tracking “conserved loops” and “regression-containing loops” in mice from P5 to P21 (Figure 3A). Interestingly, not accounting for P5 ECs fated for regression (white asterisks), by excluding them from the analysis, showed that regression-containing loops displayed a significant increase in the change of cell number compared to conserved loops (Figure 3A & 3A’). However, when regressed ECs were included in the P5 cell counts, there was no longer a significant difference in the net change of EC number between conserved loops and regression-containing loops, indicating that regressed ECs were incorporated into the P21 capillary loops (Figure 3A’’). We next investigated whether EC proliferation or apoptosis contributed to the regulation of EC density in the remodeling plexus. Phospho-Histone H3 whole-mount immunostaining showed that EC proliferation is negligible (<0.4%) during this postnatal period (Supplementary Figure 5). Furthermore, when surveying EC nuclei morphology in the plexus of neonates, we did not detect apoptotic bodies, a method we have previously used to score apoptosis events reliably (data not shown) (*32*). These data demonstrate that EC incorporation from regressed vessel segments is the main contributor to the regulation of capillary loop density during postnatal development.

**Figure 3.**
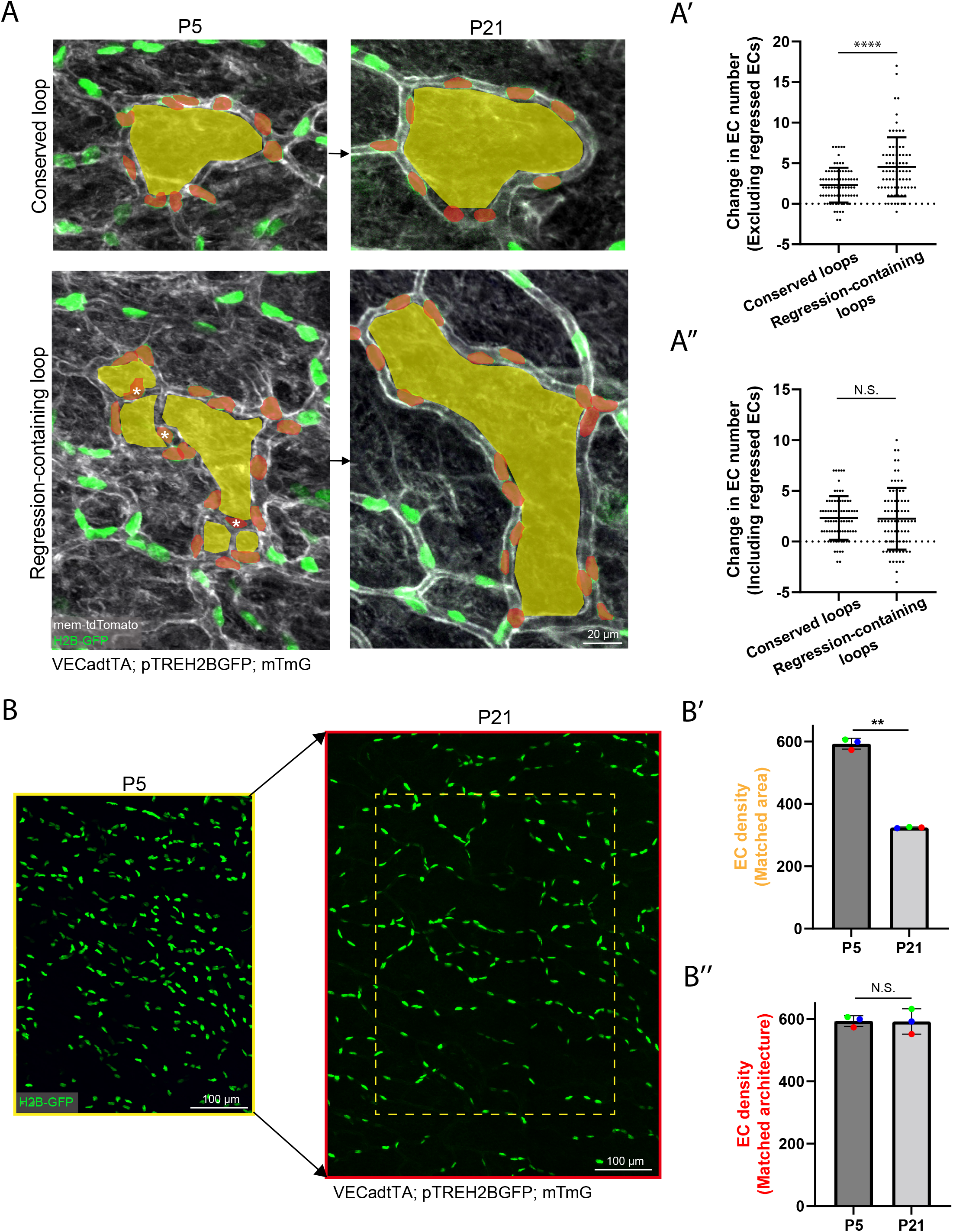
Cellular density in the developing plexus is locally controlled by the incorporation of ECs from regressed segments. (A) Comparison of changes in EC number in conserved and regression-containing loops tracked from P5 to P21 in *VECadtTA;pTREH2BGFP;mTmG* mice. Examples of conserved loops (top panel) and regression-containing loops (bottom panel) with pseudocolored loops in yellow and nuclei within loops of interest in red. ECs within regressed vessels are marked with white asterisks. (A’) Quantification of the change in EC number between conserved loops and regression-containing loops excluding regressed ECs (white asterisks) or (A’’) including regressed ECs (n=150 capillary loops from 3 mice). N.S, not significant; ****, P<0.0001 by unpaired Student’s t-test. (B) Tracking of EC nuclei within the same vascular architecture in P5 and P21 mice allows for the interrogation of how EC density adapts to plexus expansion. (B’) Quantification of EC density normalized to area shows that overall density decreases in P21 compared to P5 animals when controlled for area (yellow box indicates the matched P5 area in the P21 plexus) (n=3 mice). (B’’) Quantification of EC density matched to conserved vessel architecture (red outline indicates matched P5 architecture at P21) shows that EC density is largely preserved within the same vascular structures that have undergone expansion (n=3 mice). N.S, not significant; **, P<0.01 by unpaired Student’s t-test.

We next investigated potential plexus-wide changes in EC density. We employed our EC nuclear reporter approach and analyzed large numbers of ECs (>500 cells) during plexus expansion from P5 to P21 (Figure 3B). Interestingly, when controlling for the conserved area, EC density was reduced by approximately 2-fold in P21 compared to P5 mice (Figure 3B’). However, EC density was largely maintained when assessed relative to the original, architecturally conserved vessel structures (Figure 3B’’). These findings indicate that cellular density does not scale proportionally with plexus expansion and that tight regulatory mechanisms ensure proper EC distribution throughout the skin vasculature.

Postnatal cardiomyocytes have been shown to undergo increases in size that result in heart growth (*33*). To investigate whether a similar response is functional in the skin vasculature, we assessed whether changes in EC size contributed to loop expansion. Our experimental approach consisted of analyzing the minimum distance between EC nuclei and their closest neighbors. We found that the internuclear distance between neighboring ECs was significantly increased in P21 compared to P5 vessels, confirming that EC growth contributes to plexus expansion (Supplementary Figure 6). Together, our results demonstrate that EC density in the remodeling plexus is regulated by the incorporation of ECs from regressed vessels combined with increases in EC size. These two mechanisms allow for the expansion of the capillary network during postnatal development.

### ECs become positionally stable in adulthood but coordinate neighborhood rearrangement in response to local injury

The gradually decreasing rate of EC migration during neonatal development prompted us to investigate whether ECs ultimately cease to migrate or continue to change positions in the adult plexus, albeit at slower rates. To approach this issue, we labeled ECs in 5 week-old mice (P35) and revisited the animals 1 month later at 9 weeks of age (P63). Analyses of the amount of overlap of EC membrane occupancy in the overlay of the two time-points, and quantification of the membrane displacement of these labeled cells show negligible amounts of displacement over this 1-month interval (Figure 4A & 4A’). The observed cessation of cellular mobility parallels the lack of vessel remodeling observed in adult vessels.

**Figure 4.**
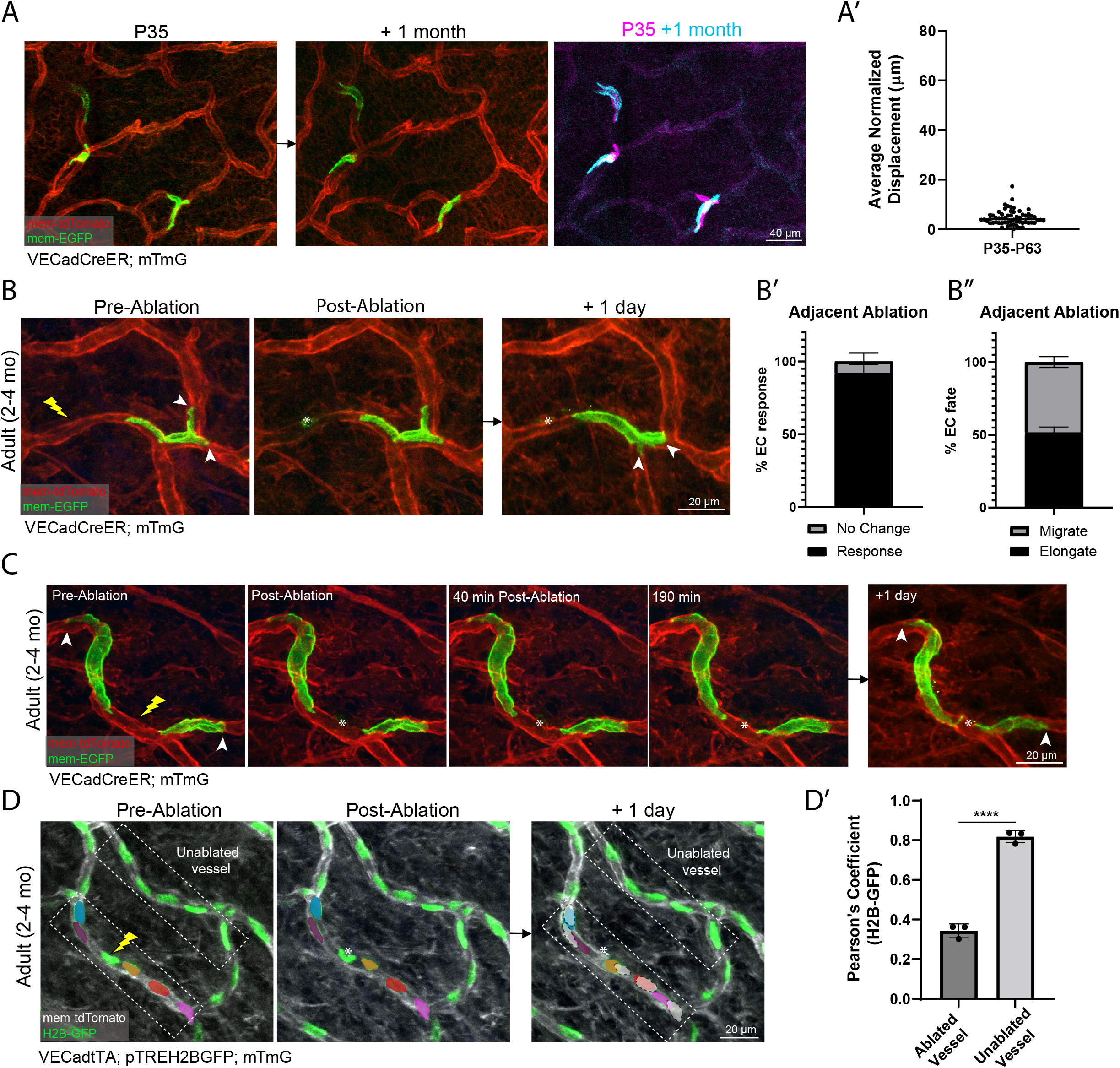
ECs become positionally stable in adulthood but coordinate neighborhood rearrangement in response to local injury. (A) Single labeled ECs in *VECadCreER;mTmG* mice at P35 longitudinally tracked over a 1 month interval are found to have undergone minimal levels of positional change. Overlay of ECs (magenta) at P35 and 1 month later (cyan) shows substantial overlap of the membrane occupancy between these timepoints. (A’) Quantification of migration between P35-P63 indicates that the majority of ECs undergo negligible levels of membrane displacement (<10 micron) in the mature plexus (n=60 cells from 3 mice). (B) Adult mice (2-4 months) inflicted with targeted laser ablation adjacent to single labeled ECs. 24 h revisit reveals that ECs are capable of reactivating their migratory capacity, releasing their distal site of anchorage and migrating towards the injury site. (B’) The vast majority (>90%) of all labeled cells respond to an adjacent laser injury. (B’’) Cells adjacent to an injury respond by migrating or elongating towards the injury site with approximately equal probability (n=180 cells from 3 mice). (C) Time-lapse imaging of a laser ablation targeted to a site in between two flanking labeled ECs, followed by 24 h revisit, shows that more than one cell is capable of responding to local injury. (D) Labeling of all EC nuclei (*VECadtTA;pTREH2BGFP;mTmG*) demonstrates the network dynamics within a capillary loop inflicted with laser injury and revisited 24 h later. EC network response was largely limited to ECs occupying the ablated vessel segment (pseudocolored nuclei) while cells in the opposing segment of the same loop appear to be unaffected. White cell masks depict the position of ECs prior to ablation. (D’) Pearson’s correlation coefficient of EC nuclei pre-ablation and 24 h post-ablation in the ablated vessel segment compared to nuclei in the opposing segment of the same capillary loop (n=3 mice). ****, P<0.0001 by unpaired Student’s t-test. For all images, lightning bolt denotes the site of laser ablation; white arrowheads denote the distal site of EC anchorage; white asterisks denote the injury site.

Considering the stability of the young adult plexus, we next sought to determine how vessel maintenance is achieved following damage or neighboring cell loss. To address this question, we induced local damage using targeted two-photon laser ablation of selected vessels. These micro-injuries were directed at sites adjacent to single labeled ECs (Figure 4B). This approach allowed for visualization and assessment of nearby responses when the injured area was revisited 24 hours later. Interestingly, we found that neighboring ECs responded to injury by either reactivating their migratory behavior and moving towards the damaged site or elongating towards the injury without relinquishing their anchorage site (Figure 4B & 4C). Quantification of the cumulative responses of injury-adjacent ECs showed that >90% of the labeled cells responded by either elongating or migrating towards the damage site with equal probability (Figure 4B’ & 4B”). We next investigated whether single or multiple neighboring ECs were involved in the repair process by inducing targeted ablation in regions containing multiple labeled ECs (Figure 4C). Using time-lapse imaging, we demonstrated that two flanking ECs respond rapidly by remodeling their leading edge and elongating towards the damage site during the first few hours following ablation (Figure 4C; Supplementary Movie 1). These cells ultimately closed the distance between each other, as observed in a 24-hour revisit (Figure 4C).

To further test the extent of neighbor participation in damage repair, we used our laser ablation approach in a model where all EC nuclei were labeled. Interestingly, we detected no EC replacement or addition in 24-hour revisits that followed laser injury. These results showed that EC proliferation is not a mechanism used by adult capillaries to repair local damage. We next determined the position of ECs before and after (*i.e*., 24 hour) injury and compared them with that of vessel segments on the opposing side of the same capillary loop. We found that the network response extended to approximately 2-3 ECs from the injury site in either direction (Figure 4D; pseudo-colored nuclei; dotted boxes); multiple neighbors participated in the repair process (Figure 4D’). Collectively, these data demonstrate that adult ECs are positionally stable under homeostatic conditions. Following injury, these cells mount a coordinated rearrangement response that effectively repairs the skin vasculature.

### Adult but not neonatal ECs preferentially activate a plasmalemmal self-repair mechanism to survive injury and preserve the vascular architecture

To better understand the repair process, we next interrogated the mechanism by which damaged ECs are cleared while maintaining the structural integrity of the vessels. We visualized the spatiotemporal dynamics of EC death and clearance following targeted laser ablation using time-lapse microscopy. Remarkably, we found that adult ECs did not die; instead, they survived by excising portions of their cell body upstream of the damage site within the first hour following ablation (Figure 5A; white arrowhead; Supplementary Movie 2). This response was followed by EC elongation towards the injury location. We next determined the fate of injured ECs 24 hours following laser ablation (Figure 5B; top panel). We found that the majority of membrane-damaged ECs (>60%) were present 24 hours following laser injury (Figure 5B & B’). In contrast, only ~20% of neonatal (*i.e*., P5-P7) ECs were maintained 24 hours post-ablation (Figure 5B & 5B’). These observations show that plasmalemmal self-repair is utilized by adult ECs to a much greater extent (*i.e*., 3-fold) compared to neonatal ECs.

**Figure 5.**
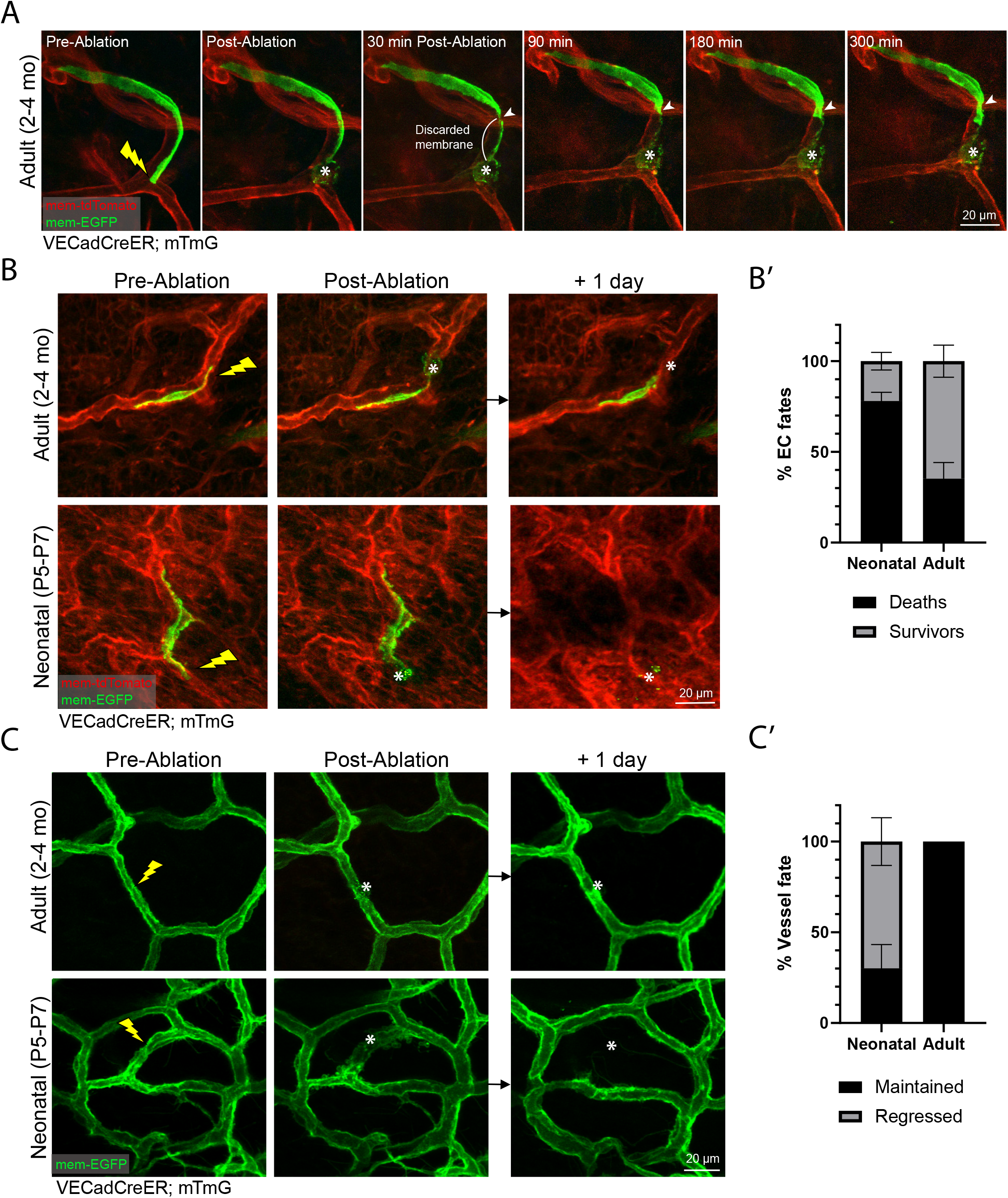
Adult but not neonatal ECs preferentially activate a plasmalemmal self-repair mechanism to survive injury and preserve the vascular architecture. (A) Time-lapse imaging of a labeled EC of a *VECadCreER;mTmG* mouse inflicted with laser damage shows that ECs are capable of carrying out plasmalemmal self-repair, excising a portion of its cell body upstream of the site of damage, before elongating back towards the injury. (B) (Top panel) An adult EC induced with laser damage followed by 24 h revisit shows that adult ECs are still viable after carrying out plasmalemmal self-repair. (Bottom panel) A neonatal EC inflicted with laser damage followed by 24 h revisit shows that neonatal ECs are eliminated following ablation. (B’) Quantification of EC fates after laser injury reveals that the majority (~60%) of adult ECs are viable following laser ablation while only 20% of neonatal ECs remain 24 h post-ablation (n=60 neonatal cells; n=84 adult cells from 3 mice respectively). (C) Maximal labeling of the vessel network allows for visualization of vessel fate following ablation. (Top panel) Adult vessel segments are maintained 24 h post-ablation. (Bottom panel) Neonatal vessel segments are often eliminated or induced to regress 24 h post-ablation. (C’) Quantification of vessel segment fates shows that while adult vessel segments are maintained 100% of the time following injury, only ~30% of neonatal vessel segments are maintained 24 h post-ablation (n=88 neonatal vessels; n=99 adult vessels from 3 mice respectively). For all images, lightning bolt denotes the site of laser ablation; white arrowheads denote the site of excision; white asterisks denote the injury site.

The observed differences in cell survival following ablation in neonatal compared to adult ECs prompted us to investigate the implications at the tissue level. We assessed the fate of vessel segments by tracking the same tissues following laser injury and showed that *all* adult vessels evaluated were repaired following laser ablation (Figure 5C & 5C’). In contrast, neonatal vessels were maintained in only ~30% of the vessels tracked; most injured neonatal vessels underwent regression following laser ablation (Figure 5C & 5C’). Together, our findings point at an age-dependent divergence in the regulation of EC membrane repair in response to localized damage. These developmental differences directly impact the manner in which the vascular architecture is maintained or remodeled.

### Neonatal vessel regression and adult vascular maintenance are orchestrated by VEGFR2 dependent signaling

Our work elucidates the process by which maturation of the vasculature is orchestrated by age-dependent differences in tissue and cellular behaviors. To understand how maturation of the vasculature is orchestrated at the molecular level, we turned to a critical vascular signaling pathway-VEGF-A signaling. While the role of the VEGF-A signaling in regulating angiogenesis during embryonic development and tumor growth is well established, much less is known in the context of developmental vessel regression and vascular network maturation (*34*). We first sought to assess the expression of VEGF-A and its cognate receptor VEGFR2, the primary downstream effector of VEGF-A mediated signaling, during postnatal skin development. Utilizing a VEGF-A-specific ELISA that recognizes the VEGF-A_164_ and VEGF-A_120_ isoforms of murine VEGF-A, we determined that VEGF-A expression was significantly higher (~3-fold) in P6 skin lysates when compared to P15 animals (Figure 6A). There was no difference in P15 versus adult skin (2 months) (Figures 6A). Whole mount immunostaining for VEGFR2 shows enriched expression in blood vessels at P5 in contrast to broad dermal expression at P21 (Figure 6B). Furthermore, FACS analysis of VEGFR2 expression on ECs reveals that it is significantly lower at P21 compared to P5 (Supplementary Figure 7A, 7B & 7B’). Altogether, analyses of ligand and receptor protein expression suggest that VEGF-A signaling via VEGFR2 may play a regulatory role during the skin vascular maturation process.

**Figure 6.**
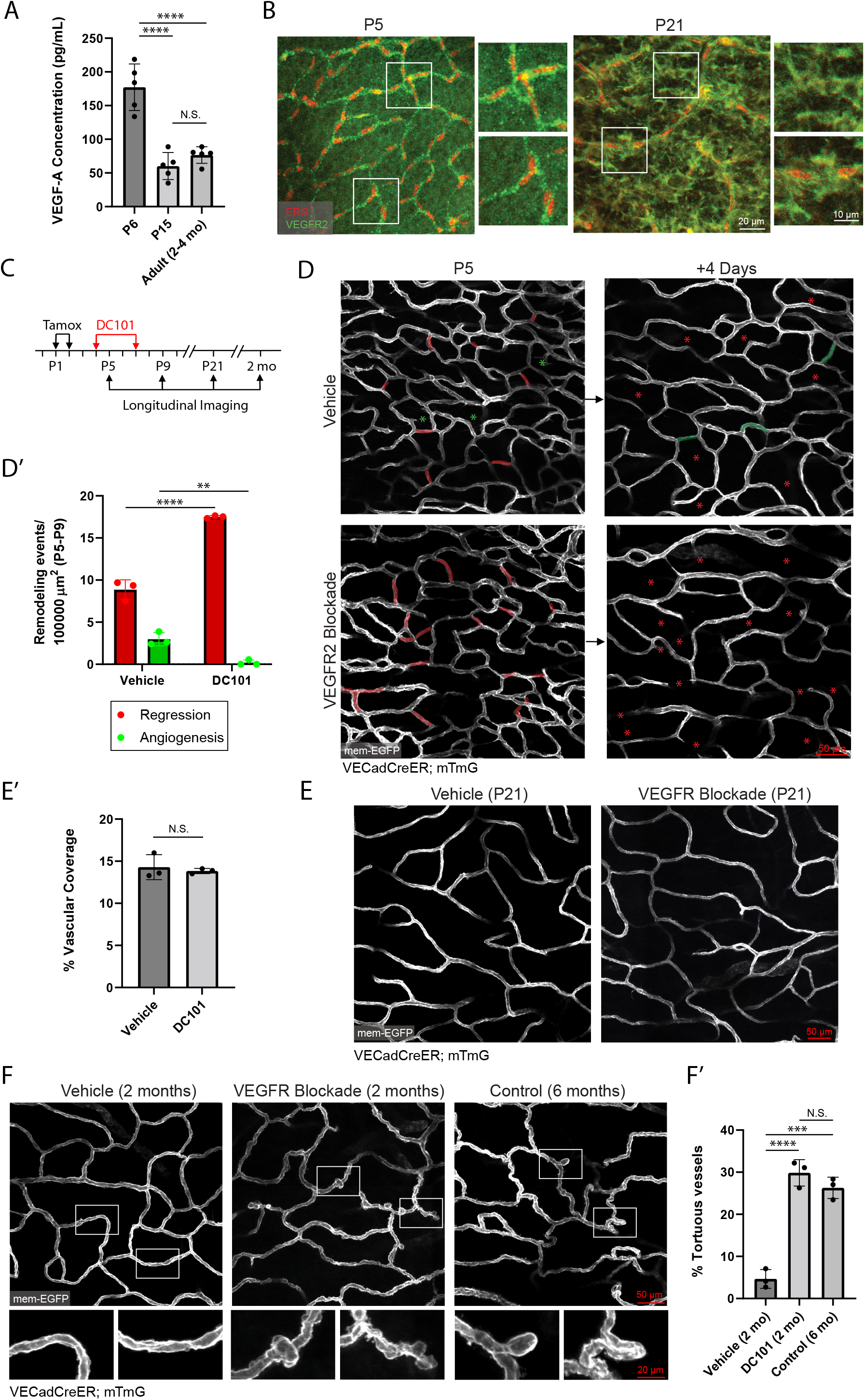
Neonatal vessel regression and adult vascular maintenance are orchestrated by VEGFR2 dependent signaling. (A) VEGF-A specific ELISA carried out on samples generated from full thickness skin shows the changes of VEGF-A expression levels between P6, P15 and adult mouse skin (n=5 mice per condition). N.S, not significant; ****, P<0.0001 by one-way ANOVA followed by Tukey’s post-hoc test. (B) Whole-mount immunostaining of VEGFR2 (green) and ERG (red) as a marker of EC nuclei in P5 and P21 mouse skin. VEGFR2 displays EC specific localization at P5, as opposed to broad expression in various dermal cells at P21. (C) Schematic of DC101 administration at P4 and P7, followed by longitudinal imaging at P5, P9, P21 and adulthood (2 months). (D & D’) Longitudinal imaging of vehicle (PBS) versus DC101 treated mice from P5 to P9 shows that DC101 treated animals display significantly higher levels of vessel regression (n=3 mice). Regressed vessels are pseudocolored in red at P5 and red asterisks at P9. New vessels from angiogenesis are pseudocolored in green at P9 with green asterisks at P5. **, P<0.01; ****, P<0.0001 by unpaired Student’s t-test (E) Comparison of the vascular plexus of vehicle and DC101 treated animals at P21. (E’) Quantification of vascular coverage in vehicle and DC101 treated animals shows no significant difference by unpaired Student’s t-test (n=3 mice). N.S, not significant. (F) Comparison of the vascular plexus of 2 month old vehicle, 2 month old DC101 treated, and 6 month old control mice shows the abnormal tortuous morphology (insets) of DC101 treated animals and 6 month old controls compared to 2 month old vehicle treated mice. (F’) Quantification of the percentage of vessels exhibiting tortuous morphology shows significantly higher levels of tortuous vessels in 2 month old DC101 treated and 6 month old control compared to 2 month old vehicle treated mice (n=3 mice). N.S, not significant; ***, P<0.001; ****, P<0.0001 by one-way ANOVA followed by Tukey’s post-hoc test.

To test whether VEGFR2 signaling plays a role towards skin vascular maturation, we employed a murine neutralizing antibody, DC101, a potent and specific inhibitor of VEGFR2 activation and downstream signaling (*35*, *36*). Administration of DC101 at P4 followed by imaging the next day showed the dissolution of vessel sprouts and lateral filopodial extensions along the vessel surface, demonstrating the efficacy of VEGF-A signaling blockade (Supplementary Figure 7C, 7C’ & 7C’’). To determine whether VEGFR2 blockade affects vessel remodeling process, we longitudinally tracked mice treated with either vehicle or DC101 (two doses at P4 and P7) from P5 to P9 (Figure 6C). DC101 treated animals displayed ~2-fold higher levels of vessel regression in the P5-P9 interval when compared to vehicle treated littermates (Figure 6D & 6D’), consistent with a previous study that utilized genetic deletion of VEGFR2 (*37*). Further tracking at P21 show overall healthy plexus morphology and vessel coverage when compared to vehicle treated littermates (Figure 6E & 6E’), suggesting that treatment led to an acceleration of the physiological program for vessel remodeling.

Lastly, to understand the longer-term consequences of neonatal VEGFR2 blockade, we continued to track these mice into adulthood at 2 months of age. Intriguingly, when compared to vehicle treated controls, DC101-treated animals displayed abnormal vessel morphology, with widespread distribution of vessel kinks and tortuous capillaries throughout the plexus, which are established indicators of unhealthy or diseased vessels (Figure 6F & 6F’) (*38*–*41*). This DC101 induced capillary morphology was reminiscent of vessel structures that we had previously observed in control 6-month old animals (Figure 6F). These results show that blockade of vascular VEGFR2 signaling during neonatal stages leads to premature deterioration of vessel structure and more aged morphology. These results emphasize the importance of the postnatal vascular maturation process and how dysregulation of VEGF-A/VEGFR2 signaling during this critical window can lead to impaired vascular homeostasis in adulthood.

## DISCUSSION

This study shows that neonatal vessel remodeling in the expanding plexus is driven primarily by network-wide vessel regression. The majority of new vessels formed via angiogenesis represent transient connections fated for regression. These findings indicate that neonatal remodeling of skin capillaries favors an overall reduction in vessel coverage and raises several questions regarding the purpose and regulation of these transient vessel connections. It was previously proposed that external signals promote the formation of new vessels fated for elimination as a strategy to favor regression of immature vessel segments (*42*). Alternatively, these vascular structures may play functional roles in the stereotyped remodeling of the developing plexus. Future studies should focus on elucidating the regulation of transient vessel formation as this process may have important implications for angiogenic remodeling in other developmental systems or pathological states.

We find that plexus-wide rearrangement and incorporation of ECs from regressed vessels serves as a mechanism to regulate EC density in the growing plexus. It is well established that ECs adopt cellular quiescence in adulthood and actively repress their proliferative capacity under homeostatic conditions (*43*, *44*). Our findings suggest that the suppression of EC proliferation begins early in postnatal life, as the capillary plexus favors the reabsorption of regressed ECs and increases in EC size over increasing EC number as a mechanism for plexus expansion. These results shed light upon shared and divergent cellular mechanisms that underlie vessel regression in developmental versus pathological states (*42*). This could advance our understanding of pathologies or states that involve capillary rarefaction, defined as a reduction of capillary density, with particular relevance to conditions that afflict the skin vasculature, such as scleroderma, primary hypertension, and aging (*18*–*20*).

Establishing the positional and architectural stability of ECs during adulthood allowed us to investigate the cellular mechanisms utilized by ECs to maintain homeostasis. We observed that capillary ECs do not proliferate in response to ablations inflicted on neighboring cells; instead, they elongate or migrate towards the injury site, suggesting that ECs compensate for the loss of neighbors through membrane extension. These results are thought-provoking in light of recent studies reporting that tissue resident ECs in large-caliber heart and liver vessels proliferate in response to physical denudation or acute genotoxic ablation (*45*, *46*). The observations raise the question of whether there is a threshold of capillary EC loss that leads to proliferative repair and to what extent angiogenesis or new vessel formation might be required for homeostatic maintenance (*41*, *47*, *48*). Subsequent investigations should also focus on differences or similarities of repair mechanisms in vessels of differing caliber.

In addition to neighboring EC responses, we uncovered a divergence of fates in neonatal versus adult ECs following ablative injury. The ability of adult ECs to self-repair is reminiscent of the membrane resealing capacity observed in muscle fibers, an ability believed to have evolved owing to the frequency of membrane injuries resulting from repeated muscle contraction (*49*, *50*). Considering the persistent exposure of ECs to shear and contractile forces associated with blood flow, it is plausible that ECs evolved similar capacities for self-repair as a protective measure against injuries related to these forces.

Interrogation of the molecular regulation of skin vascular maturation showed that VEGF-A expression levels are downregulated in adulthood, along with a decrease in EC specific VEGFR2 expression. Our finding that acute neonatal blockade of VEGFR2 signaling is sufficient to accelerate the vessel regression program indicates that VEGFR2 is a critical regulator of vessel maturation status. Future work should focus upon the role of pericytes in regulation of vessel regression and its role in VEGFR2 dependent vessel stabilization, as previous studies have found that VEGF-A/VEGFR2 signaling is antagonistic to pericyte recruitment during neovascularization (*51*). Our results also suggest that proper vascular network maturation depends upon a gradual downregulation of VEGF-A receptor and/or ligand expression levels at a specific time window during neonatal stages, the failure of which leads to eventual deterioration of vessel morphology and health in adulthood (Figure 6).This finding highlights the importance of the neonatal developmental window as it has been shown that removal of VEGF-A blockade in adult mice can lead to a restoration of normal vessel architecture. Subsequent investigation should center on understanding the mechanism by which acute inhibition of VEGFR2 signaling in early life can lead to deterioration of vascular homeostasis.

In conclusion, our work elucidates novel aspects of the developmental journey of the skin capillary network and identifies the principles that guide the remodeling of this network as it grows proportionately with the organism at the tissue and cellular levels to achieve adult homeostasis (Figure 7). We identified a long-lived positional stability adopted by ECs during adult homeostasis and unveiled strategies employed by network ECs to maintain vessel integrity in adulthood. We discovered that adult ECs possess the remarkable capacity to self-repair and maintain vessel architecture, features not exhibited to the same extent by neonatal ECs. Lastly, we found that acute blockade of VEGF signaling in early life accelerates the neonatal remodeling program leading to adverse consequences to vessel architecture in adulthood. Future studies will need to relate these tissue and cellular mechanisms to blood flow by developing imaging modalities that simultaneously assess hemodynamic parameters. This issue is important in light of previous studies reporting an inverse relationship between the directionality of EC migration and blood flow (*52*–*54*). In summary, our work describes the postnatal journey of ECs and their role in the maturation and maintenance of the capillary network. It establishes a new understanding of the cellular mechanisms that dictate neonatal vascular development and adult homeostasis.

**Figure 7.**
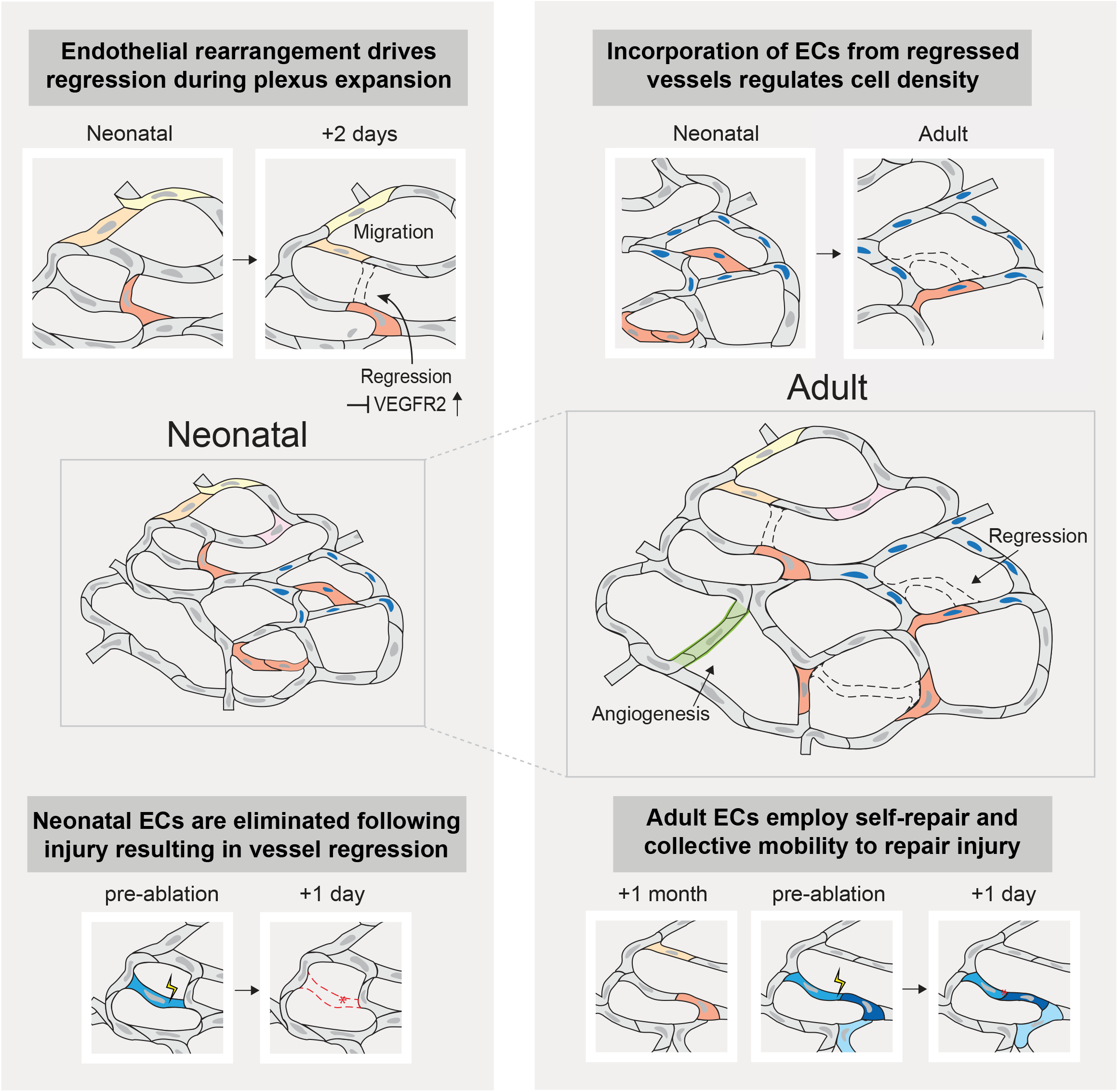
Schematic representation of EC behaviors during neonatal vs adult stages. During neonatal development, ECs rearrange their positions and execute vessel regression through migration and incorporation into adjacent vessels. This rearrangement and addition of cell numbers through regression serves as a mechanism to regulate EC density during plexus expansion. In adulthood, ECs become positionally stable. In response to vessel injury, adult ECs are able to reengage migration and execute plasmalemmal self-repair to survive damage and maintain vessel integrity. In contrast, neonatal ECs are disposed towards death and induced vessel regression following membrane injury.

## MATERIALS & METHODS

### Mice

*VE-CadherinCreER (VECadCreER*) mice (*24*) were obtained from Ralf Adams (Max Planck Institute). Elaine Fuchs (Rockefeller University) generously provided *pTRE-H2BGFP* mice (*31*). *Rosa26-loxP-membraneTomato-(stop)-membraneGFP (mTmG) (25*) and *VE-CadherintTA (VECadtTA) (30*) mouse lines were from The Jackson Laboratory. All mice were bred to a mixed CD1 albino background. The animals used in this study were of both genders and within an age range of P1 to P21 for postnatal development studies and 2-4 months for adult studies. All experimentation involving animal subjects was approved by the Institutional Animal Care and Use Committee at Yale School of Medicine and conducted following the approved animal handling protocol.

### In vivo imaging

In vivo imaging procedures were carried out as previously described (*17*). Mice were anesthetized in an isoflurane chamber and continuously received vaporized isoflurane through a nose cone (1% oxygen and air) on a warm heating pad for the duration of the analysis. A LaVision TriM Scope II (LaVision Biotec) laser scanning microscope equipped with a Chameleon Vision II (Coherent) two-photon laser (using 940 nm for live imaging and whole-mounts) and a Chameleon Discovery (Coherent) two-photon laser (using 1120 nm for live imaging and whole-mounts) was used to acquire z-stacks of 100–200 μm in 1–2 μm steps through either a Nikon 25x/1.0 or a Nikon 40x/1.15 water immersion objective. Optical sections were scanned with a field of view of 0.44 x 0.44 mm^2^ or 0.28 x 0.28 mm^2^, respectively, at 700 Hz. To visualize large areas, we imaged 4-24 tiles of optical fields with a motorized stage and automatically acquired sequential fields of view, as previously described. For time-lapse imaging, serial optical sections were obtained in a range of 5-20 min intervals, dependent upon experimental setup with a duration of 1-5 hours.

### Tamoxifen induction & VEGFR2 neutralization

To induce maximal expression of membrane-GFP labeling in neonates, *VECadherin-CreER; mTmG* mice received intragastric injections of 100 μg of tamoxifen at postnatal days 1 (P1) and P2. Tamoxifen (Sigma) stocks were prepared by dissolving 200 mg of tamoxifen powder in 10 mL of corn oil. 2 mg aliquots were stored at −20°C and diluted in the required volume of corn oil before injection. Inhibition of VEGFR2 was carried out by intraperitoneal injection of an anti-VEGFR2 neutralizing antibody (DC101, BioXcell) at a concentration of 20 mg/kg in PBS or vehicle (PBS) alone. DC101 or vehicle treatment was carried out at P4 and P7.

### Laser ablations

Laser ablations were carried out using a Chameleon Vision II (Coherent) two-photon laser at a wavelength of 810 nm, using either a Nikon 25x/1.0 or Nikon 40x/1.15 water immersion objective with 10% laser power. Experimentally, the field of view of the scanning head was reduced to a 1 μm^2^ sized window and positioned at the location to be ablated. We then activated the “live” scan function until a mild autofluorescence signal became visible in the green channel (~3-5 seconds). For adjacent injury experiments, ablations were positioned ~30 μm away from a labeled EC. For experiments involving ablation of the labeled EC, ablations were carried out on the tip of EC membranes to ensure consistency of ablations from cell to cell and maximize the distance between the site of ablation and the nucleus as nuclear ablation invariably leads to cell death.

### Image analysis

Raw image stacks were imported for analysis into FIJI (ImageJ v1.52p, NIH) or Imaris software (v.9.5.1; Bitplane/Oxford Instruments). Individual optical planes, summed or max Z stacks of sequential optical sections were used to assemble figures. The tiled images were stitched by a grid/collection stitching plugin in Fiji. Prism software (Graphpad, v.8.0.0) was used to graph the data. Average normalized displacement was determined based on the schematic and calculations depicted in Supplementary Figure 5. Briefly, we measured and averaged trailing- and leading-edge membrane displacements between successive time points to assess average displacement. This value was then normalized to the expansion factor of the occupied vessel to correct for vessel growth, resulting in the average normalized displacement value. Pearson’s correlation coefficients were calculated using ImageJ software.

### Whole-mount immunostaining

Mouse hind paw tissues were processed for whole-mount staining. Briefly, full-thickness paw skin was dissected and fixed in 4% paraformaldehyde in PBS for 4-6 hours at room temperature, washed in PBS, and then blocked with 0.2% Triton X-100, 5% normal donkey serum, 1 % BSA in PBS overnight at 37°C. The samples were then incubated with primary antibodies for 48-72 h and secondary antibodies for 24 h on a rocker at 37°C. The anti-phospho-histone H3 (Ser10) antibody was from EMD Millipore (06-570) and was used at a 1:100-fold dilution; anti-Flk-1 (VEGFR2) antibody was from BD Biosciences (555307) and was used at 1:100-fold dilution; anti-ERG antibody was from Abcam (AB92513) and was used at 1:200-fold dilution; goat anti-rabbit IgG H&L (Alexa Fluor®) 568 & goat anti-rat IgG H&L (Alexa Fluor®) 568 was from ThermoFisher and was used at a 1:400-fold dilution. Tissue samples were mounted on individual slides and imaged on a LaVision TriM Scope II, as described in ‘In vivo imaging.’

### Flow cytometry

P5 and P21 *VECadCreER; mTmG* mice were euthanized for fluorescence activated cell sorting (FACS) 3 days post tamoxifen induction (P2, 100 μg intragastric; P18, 2 mg intraperitoneal). Dermal single cell suspensions were prepared for flow cytometry with a protocol adapted from previously described (*55*). Briefly, back skin was incubated for 1 hour at 37 °C in 0.3 % trypsin (Sigma-Aldrich) in 150 mM NaCl, 0.5 mM KCl and 0.5 mM glucose. The dermis was physically separated from the epidermis, minced, and incubated in 2 mLs of collagenase (Sigma; C2674-11MG) at 125 U/mL final concentration in Hank’s Balanced Salt Solution (HBSS; Gibco 14170-112) with no calcium, magnesium chloride, or magnesium sulfate for 2 hours at 37°C. The resulting cells were crushed and filtered through a 70 μm filter and spun at 240G for 10 minutes. All samples were pre-treated with rat serum (Sigma-Aldrich), and incubated with anti-mouse/human CD309 (Flk-1/VEGFR2)-APC (Thermofisher, 17-5821-81; 1:100) for 30 minutes at 4°C. Samples were run on a Becton Dickinson LSRII outfitted with Diva software v 8.0.1, and the data was analyzed using Flowjo v10.6.2.

### VEGF-A Quantikine ELISA & skin sample preparation

Paw skin was excised, flash frozen with liquid nitrogen and stored at −80°C until all samples were collected. The tissue was then processed for lysate using a standard western blot protocol. In short, samples were homogenized with 100uls of RIPA buffer (Sigma-Aldrich; R0278) containing a phosphatase inhibitor (Sigma; 4906845001) and protease inhibitor (Sigma; 11697498001), incubated on ice for 30 minutes, and then spun at 15,000G for 10 minutes. Supernatants were then transferred to a clean 1.5 ml Eppendorf and stored at −80°C.

Full thickness paw tissue lysate concentration from either P6, P15 or adult mice (2 months) were first quantified using a Pierce BCA Protein Assay Kit (Thermofisher; 23225) with absorbance read using a Glomax Explorer Microplate reader (Promega) at 560nm and normalized for amount of total protein across all samples. The levels of mouse VEGF-A in paw lysates were then analyzed using a Quantikine ELISA kit specific for VEGF-A_164_ and VEGF-A_120_ (R&D Systems; MMV00) according to the manufacturer’s protocol and absorbance was read using a Glomax Explorer Microplate Reader at 450 nm.

### Statistics & reproducibility

Statistical calculations were performed using Prism software (GraphPad, v.8.0.0). A two-sided unpaired t-test was used to determine the statistical significance between two experimental conditions. One-way ANOVA followed by Tukey’s multiple comparisons test was utilized for multiple comparisons. A p-value of <0.05 was considered significant; specific p-values are noted in the figure legends. No statistical method was used to predetermine sample size (n). Mouse numbers represent the biological replicates; sample size and replicates are indicated in the figure legends. All experiments were conducted at least three times.

## ACKNOWLEDGMENTS

We thank Stefania Nicoli for providing helpful advice on the manuscript, Elaine Fuchs for sharing *pTRE-H2BGFP* mice, and Ralf Adams for supplying the *VE-CadherinCreER* mouse line.

## Author contributions

CK, KKH and VG conceived the project and designed experiments. CK, VG, and KKH wrote the manuscript. CK performed and or participated to all experiments and analyzed the data. IDS performed density analyses, whole mount experiments, and provided critical feedback to the manuscript and schematics. CMM performed VEGF-A/VEGFR2 expression analyses. PS, GS, and JB performed molecular investigations. EDM provided an initial data set for vascular exploration.

## Data and materials availability

All data from this study are available from the authors on request.

---

**Supplementary Figure 1.**
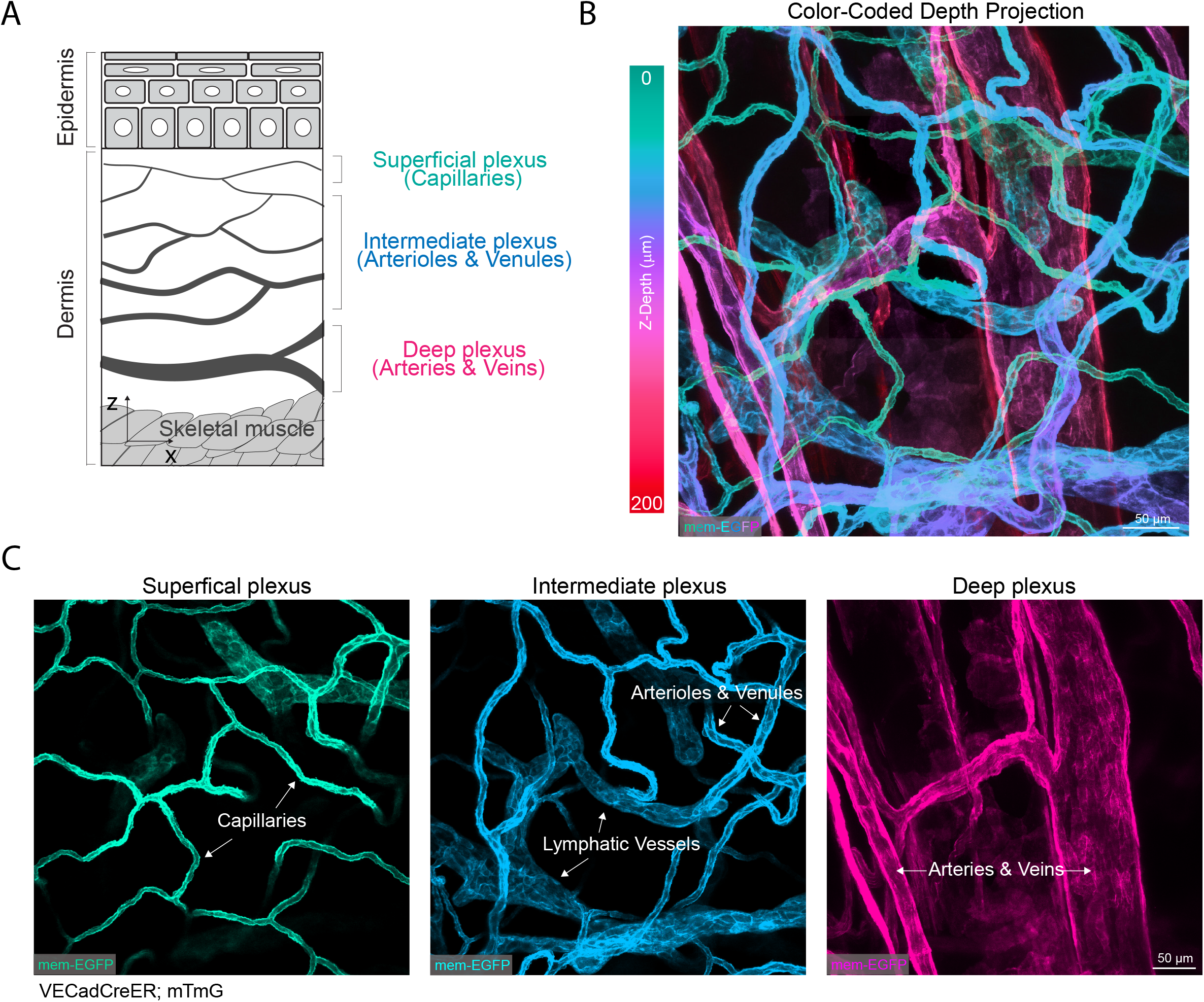
Anatomy of the murine palmoplantar skin vascular network. (A) Cross sectional schematic of the skin vascular network. Skin vasculature resides in the dermis layer of the skin beneath the barrier forming epithelial cells of the stratified epidermis. (B) Color-coded depth projection of the skin vascular network. Fully recombined *VECad-CreER;mTmG* was used to specifically label the vasculature of adult mice. Z-Depth color code on the left designates the gradient from 0 to 200 μm below the epidermis. (C) Z-projections of the superficial, intermediate, and deep plexuses.

**Supplementary Figure 2.**
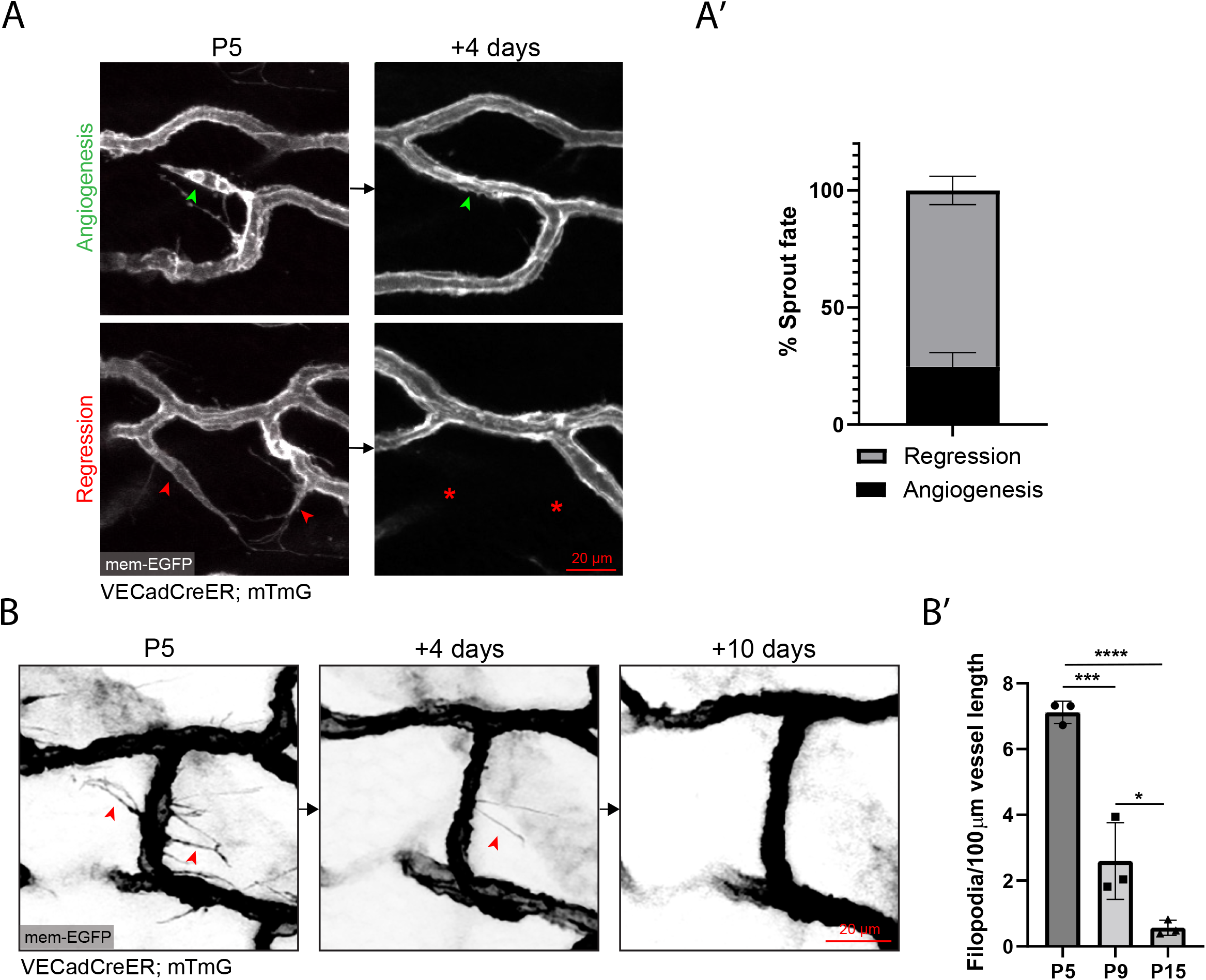
Dynamics of vascular “sprouts” and lateral filopodia in the remodeling plexus. (A) Longitudinal tracking of “sprout-like” structures from P5-P9 showing examples of angiogenesis (Green arrowheads) and regression (Red arrowheads). (A’) Quantification of sprout fates shows a significantly higher level of regression compared to angiogenesis (n=195 events from 3 mice). (B) Longitudinal tracking of lateral filopodia from P5-P9-P15 shows a reduction in these structures as postnatal development progresses. (B”) Quantification of lateral filopodia (Red arrowheads) along the vessel surface shows a significant reduction in these structures across developmental time (n=3 mice). *, P<0.05; ***, P<0.001; ****, P<0.0001 by one-way ANOVA followed by Tukey’s post-hoc test.

**Supplementary Figure 3.**
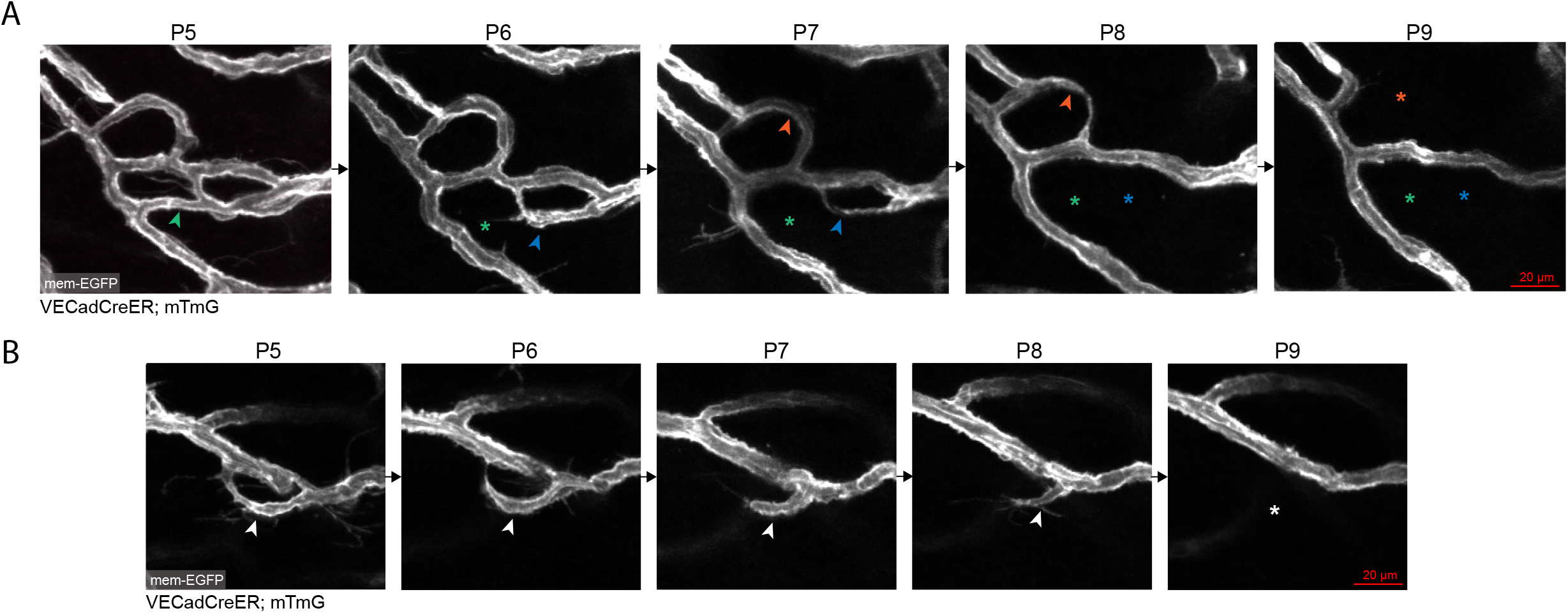
Spectrum of kinetics in the execution of vessel regression. **(A)** Daily tracking of a cluster of capillary loops undergoing sequential regression of vessel segments. Regression can be completed within a single day (Green) or 2-3 day periods (Blue & Orange). **(B)** Daily tracking of a single vessel undergoing regression depicting the progressive retraction of the regressed vessel over a period of 4 days. Arrowheads indicate a segment undergoing regression; asterisks indicate previous location of regressed segment.

**Supplementary Figure 4.**
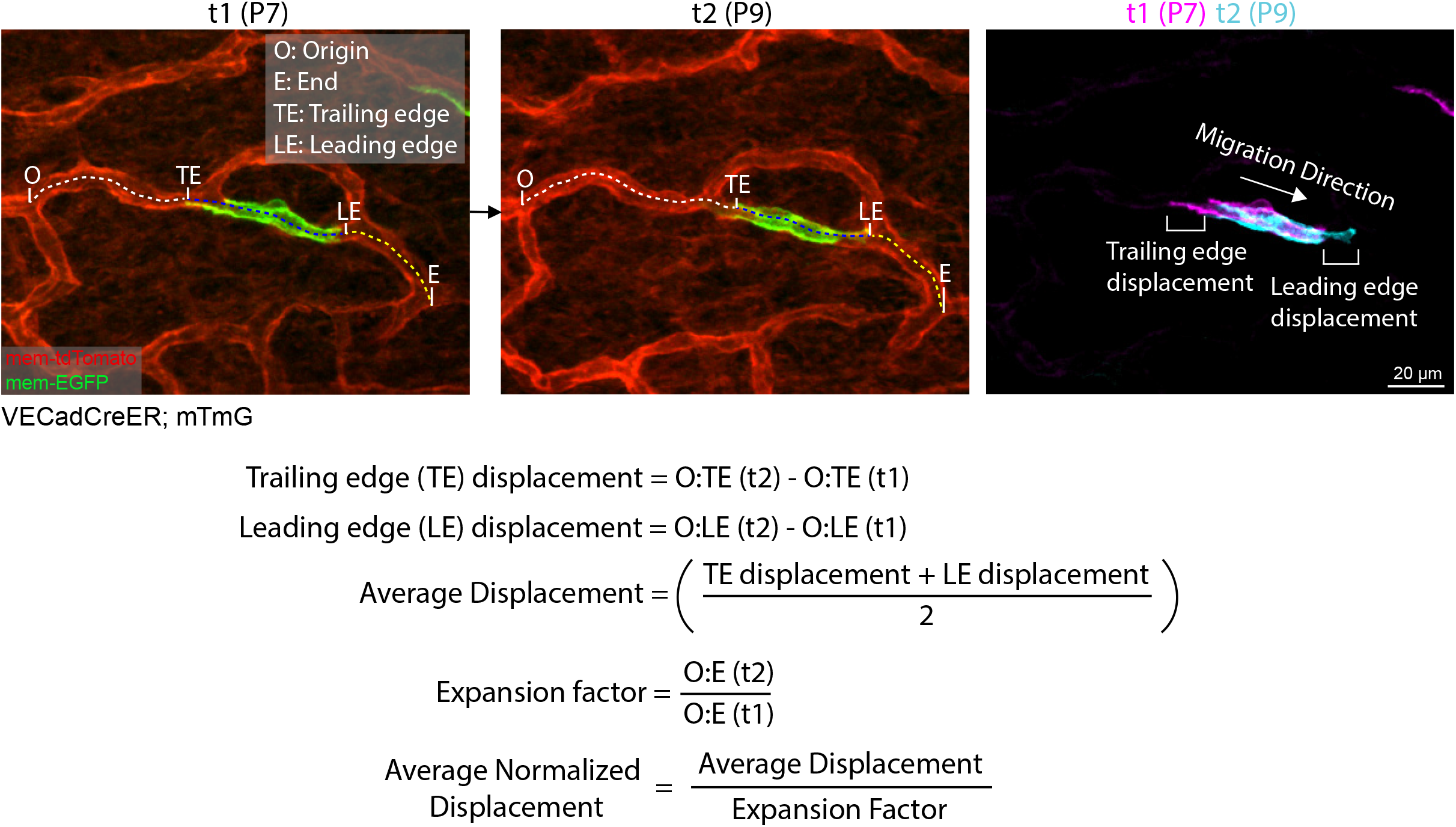
Quantification method for EC average normalized displacement as a measure of EC migration. Average displacement was quantified by averaging the change in membrane occupancy at the leading edge (LE) and trailing edge (TE) of the labeled EC between tracked time-points (t). This value was then adjusted for the expansion of the occupied segment (O:E) by normalizing the average displacement value to the expansion factor.

**Supplementary Figure 5.**
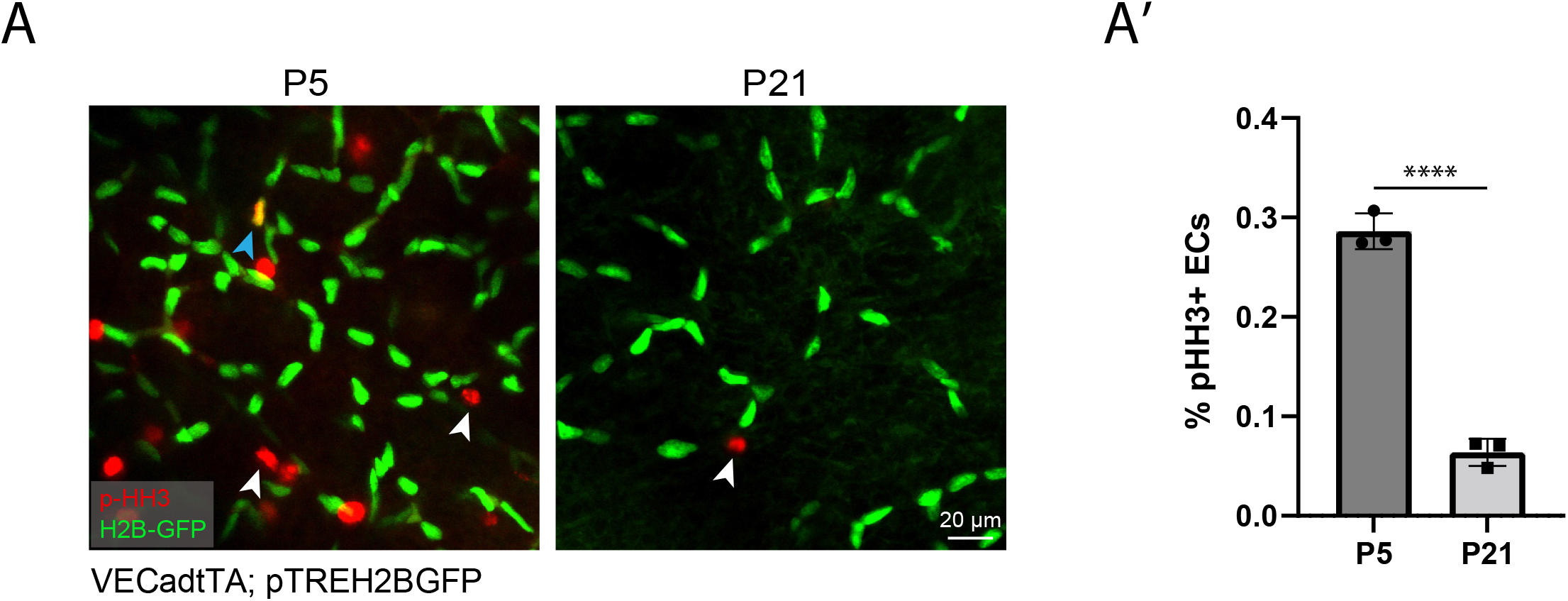
Negligible levels of EC proliferation during postnatal development. (A) Whole-mount immunostaining of phospho-Histone H3 in capillaries from *VECadtTA;pTREH2BG-FP* mouse skin at P5 and P21 ages. Proliferating non-endothelial dermal cells are indicated by white arrowheads (red nuclei) and proliferating capillary ECs are indicated by blue arrowheads (double-positive for red/green i.e yellow). (A’) Quantification of proliferating ECs shows negligible levels of proliferation (<0.4%) at P5 and P21 (n=3 mice). ****, P<0.0001 by unpaired Student’s t-test.

**Supplementary Figure 6.**
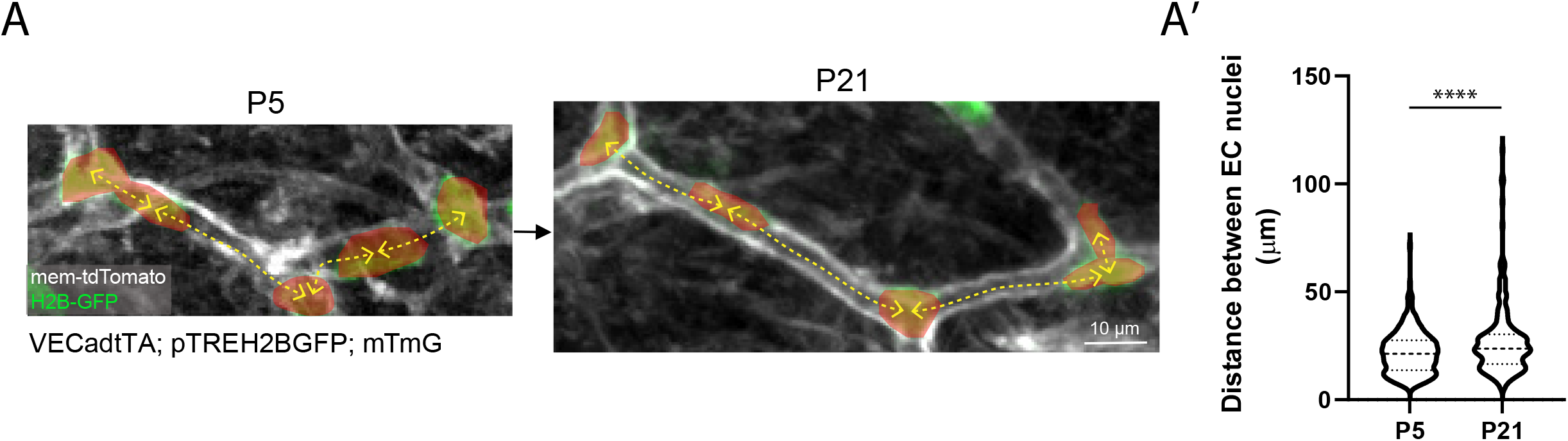
EC internuclei distance increases in the mature vasculature. (A) Schematic depicting the measurement of the minimum distance from the midpoint of EC nuclei to each of their nearest neighbors in P5 and P21 mice. (A’) Quantification of EC internuclei distance shows a significant increase in the average distance between EC nuclei in P21 compared to P5 vessels (n=233 measurements from 3 mice). ****, P<0.0001 by unpaired Student’s t-test.

**Supplementary Figure 7.**
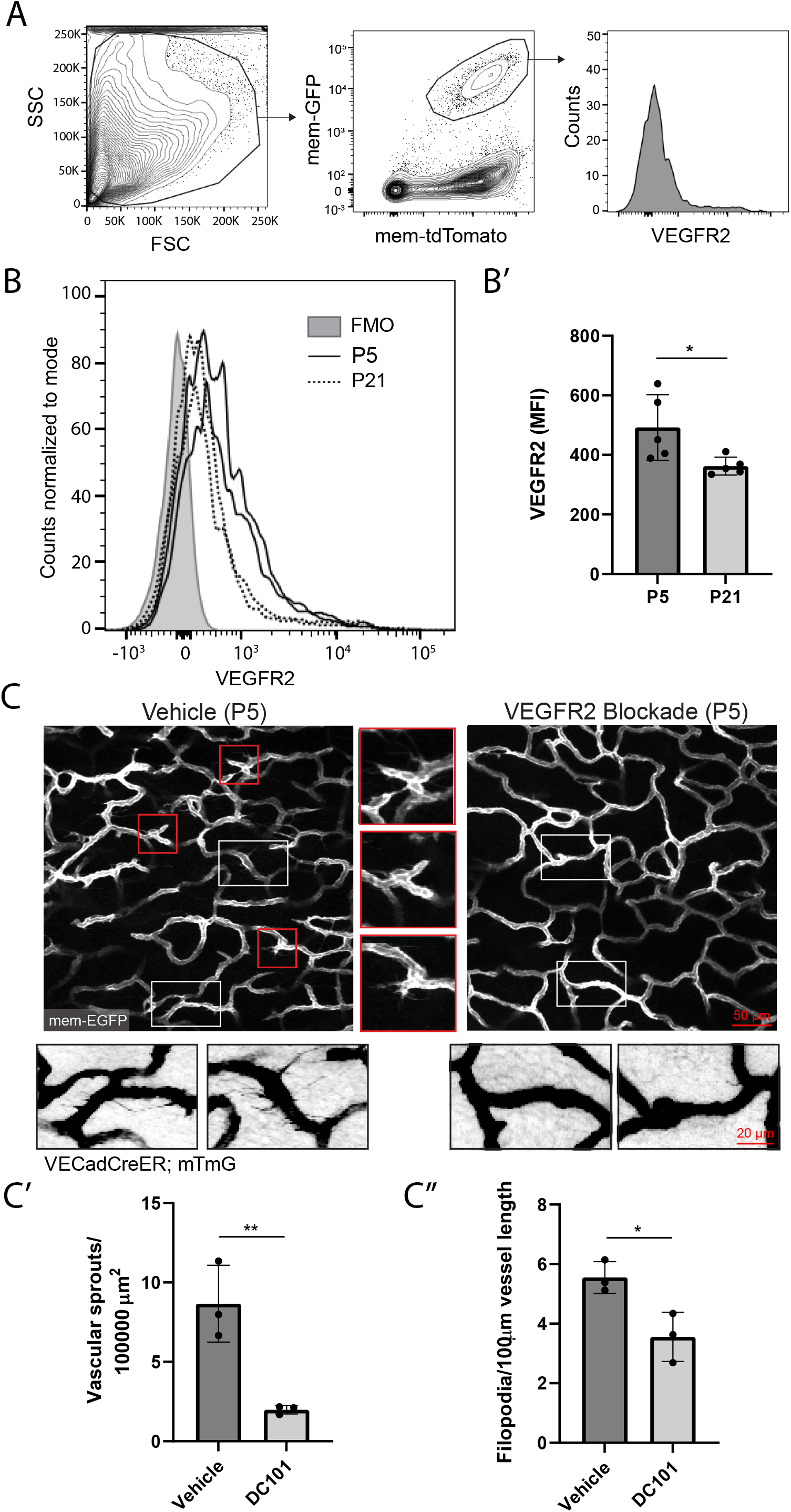
EC VEGFR2 expression decreases postnatally and is neutralized by DC101 administration. (A) Endothelial single cell suspensions were processed from the back skin of P5 and P21 *VECadCreER;mT-mG* mice 3 days post induction with tamoxifen for flow cytometry of VEGFR2 expression. Cells were first gated on FSC and SSC to include all dermal cells, and then gated on mem-GFP+ mem-tdTom+ cells to select for ECs. (B & B’) Expression levels of VEGFR2 was significantly lower in P21 compared to P5 mouse skin ECs (n=5 mice per condition; 2 representative biological replicates). *, P<0.05 by unpaired Student’s t-test. FMO was used as a control. (C) Comparison of the vascular plexus of P5 mice 24 h post administration of vehicle or DC101. Vascular sprouts (red insets) and lateral filopodia (white insets) observed in vehicle treated mice are not detectable in DC101 treated littermates. (C’ & C’’) Quantification of vascular sprouts and lateral filopodia show a significant reduction in both parameters in DC101 treated mice compared to vehicle treated littermates (n=3 mice). *, P<0.05; **, P<0.01 by unpaired Student’s t-test.

## REFERENCES

1. H. G. Augustin, G. Y. Koh, Organotypic vasculature: From descriptive heterogeneity to functional pathophysiology. Science 357, (2017).

2. M. R. Swift, B. M. Weinstein, Arterial-venous specification during development. Circ Res 104, 576–588 (2009).

3. S. F. Rocha, R. H. Adams, Molecular differentiation and specialization of vascular beds. Angiogenesis 12, 139–147 (2009).

4. C. G. Ellis, J. Jagger, M. Sharpe, The microcirculation as a functional system. Crit Care 9 Suppl 4, S3–8 (2005).

5. S. P. Herbert, D. Y. Stainier, Molecular control of endothelial cell behaviour during blood vessel morphogenesis. Nat Rev Mol Cell Biol 12, 551–564 (2011).

6. V. L. Bautch, K. M. Caron, Blood and lymphatic vessel formation. Cold Spring Harb Perspect Biol 7, a008268 (2015).

7. S. Dekoninck et al., Defining the Design Principles of Skin Epidermis Postnatal Growth. Cell 181, 604–620 e622 (2020).

8. V. Coelho-Santos, A. A. Berthiaume, S. Ornelas, H. Stuhlmann, A. Y. Shih, Imaging the construction of capillary networks in the neonatal mouse brain. Proc Natl Acad Sci U S A 118, (2021).

9. M. Takeo et al., Wnt activation in nail epithelium couples nail growth to digit regeneration. Nature 499, 228–232 (2013).

10. A. V. Gore, K. Monzo, Y. R. Cha, W. Pan, B. M. Weinstein, Vascular development in the zebrafish. Cold Spring Harb Perspect Med 2, a006684 (2012).

11. R. S. Udan, J. C. Culver, M. E. Dickinson, Understanding vascular development. Wiley Interdiscip Rev Dev Biol 2, 327–346 (2013).

12. D. Tsuruta, K. J. Green, S. Getsios, J. C. Jones, The barrier function of skin: how to keep a tight lid on water loss. Trends Cell Biol 12, 355–357 (2002).

13. M. Yokouchi et al., Epidermal cell turnover across tight junctions based on Kelvin’s tetrakaidecahedron cell shape. Elife 5, (2016).

14. N. Charkoudian, Skin blood flow in adult human thermoregulation: how it works, when it does not, and why. Mayo Clin Proc 78, 603–612 (2003).

15. T. S. Kupper, R. C. Fuhlbrigge, Immune surveillance in the skin: mechanisms and clinical consequences. Nat Rev Immunol 4, 211–222 (2004).

16. K. N. Li et al., Skin vasculature and hair follicle cross-talking associated with stem cell activation and tissue homeostasis. Elife 8, (2019).

17. C. M. Pineda et al., Intravital imaging of hair follicle regeneration in the mouse. Nat Protoc 10, 1116–1130 (2015).

18. M. Trojanowska, Cellular and molecular aspects of vascular dysfunction in systemic sclerosis. Nat Rev Rheumatol 6, 453–460 (2010).

19. T. F. Antonios, D. R. Singer, N. D. Markandu, P. S. Mortimer, G. A. MacGregor, Structural skin capillary rarefaction in essential hypertension. Hypertension 33, 998–1001 (1999).

20. L. Li et al., Age-related changes of the cutaneous microcirculation in vivo. Gerontology 52, 142–153 (2006).

21. J. D. Martin, G. Seano, R. K. Jain, Normalizing Function of Tumor Vessels: Progress, Opportunities, and Challenges. Annu Rev Physiol 81, 505–534 (2019).

22. E. Marsh, D. G. Gonzalez, E. A. Lathrop, J. Boucher, V. Greco, Positional Stability and Membrane Occupancy Define Skin Fibroblast Homeostasis In Vivo. Cell 175, 1620–1633 e1613 (2018).

23. Y. G. Kamberov et al., A genetic basis of variation in eccrine sweat gland and hair follicle density. Proc Natl Acad Sci U S A 112, 9932–9937 (2015).

24. I. Sörensen, R. H. Adams, A. Gossler, DLL1-mediated Notch activation regulates endothelial identity in mouse fetal arteries. Blood 113, 5680–5688 (2009).

25. M. D. Muzumdar, B. Tasic, K. Miyamichi, L. Li, L. Luo, A global double-fluorescent Cre reporter mouse. Genesis 45, 593–605 (2007).

26. C. A. Franco et al., Dynamic endothelial cell rearrangements drive developmental vessel regression. PLoS Biol 13, e1002125 (2015).

27. S. Abraham et al., A Rac/Cdc42 exchange factor complex promotes formation of lateral filopodia and blood vessel lumen morphogenesis. Nat Commun 6, 7286 (2015).

28. Q. Chen et al., Haemodynamics-driven developmental pruning of brain vasculature in zebrafish. PLoS Biol 10, e1001374 (2012).

29. A. Lenard et al., Endothelial cell self-fusion during vascular pruning. PLoS Biol 13, e1002126 (2015).

30. J. F. Sun et al., Microvascular patterning is controlled by fine-tuning the Akt signal. Proc Natl Acad Sci U S A 102, 128–133 (2005).

31. T. Tumbar et al., Defining the epithelial stem cell niche in skin. Science 303, 359–363 (2004).

32. K. R. Mesa et al., Niche-induced cell death and epithelial phagocytosis regulate hair follicle stem cell pool. Nature 522, 94–97 (2015).

33. M. Maillet, J. H. van Berlo, J. D. Molkentin, Molecular basis of physiological heart growth: fundamental concepts and new players. Nat Rev Mol Cell Biol 14, 38–48 (2013).

34. M. Lohela, M. Bry, T. Tammela, K. Alitalo, VEGFs and receptors involved in angiogenesis versus lymphangiogenesis. Curr Opin Cell Biol 21, 154–165 (2009).

35. M. Prewett et al., Antivascular endothelial growth factor receptor (fetal liver kinase 1) monoclonal antibody inhibits tumor angiogenesis and growth of several mouse and human tumors. Cancer Res 59, 5209–5218 (1999).

36. L. Witte et al., Monoclonal antibodies targeting the VEGF receptor-2 (Flk1/KDR) as an anti-angiogenic therapeutic strategy. Cancer Metastasis Rev 17, 155–161 (1998).

37. S. Karaman et al., Interplay of vascular endothelial growth factor receptors in organ-specific vessel maintenance. J Exp Med 219, (2022).

38. H. C. Han, Twisted blood vessels: symptoms, etiology and biomechanical mechanisms. J Vasc Res 49, 185–197 (2012).

39. C. G. Owen et al., Diabetes and the tortuosity of vessels of the bulbar conjunctiva. Ophthalmology 115, e27–32 (2008).

40. J. A. Nagy, S. H. Chang, A. M. Dvorak, H. F. Dvorak, Why are tumour blood vessels abnormal and why is it important to know? Br J Cancer 100, 865–869 (2009).

41. D. C. Chong, Z. Yu, H. E. Brighton, J. E. Bear, V. L. Bautch, Tortuous Microvessels Contribute to Wound Healing via Sprouting Angiogenesis. Arterioscler Thromb Vasc Biol 37, 1903–1912 (2017).

42. C. Korn, H. G. Augustin, Mechanisms of Vessel Pruning and Regression. Dev Cell 34, 5–17 (2015).

43. M. Ehling, S. Adams, R. Benedito, R. H. Adams, Notch controls retinal blood vessel maturation and quiescence. Development 140, 3051–3061 (2013).

44. J. Andrade et al., Control of endothelial quiescence by FOXO-regulated metabolites. Nat Cell Biol 23, 413–423 (2021).

45. A. I. McDonald et al., Endothelial Regeneration of Large Vessels Is a Biphasic Process Driven by Local Cells with Distinct Proliferative Capacities. Cell Stem Cell 23, 210–225 e216 (2018).

46. T. Wakabayashi et al., CD157 Marks Tissue-Resident Endothelial Stem Cells with Homeostatic and Regenerative Properties. Cell Stem Cell 22, 384–397 e386 (2018).

47. J. M. Arpino et al., Four-Dimensional Microvascular Analysis Reveals That Regenerative Angiogenesis in Ischemic Muscle Produces a Flawed Microcirculation. Circ Res 120, 1453–1465 (2017).

48. C. E. Evans, M. L. Iruela-Arispe, Y. Y. Zhao, Mechanisms of Endothelial Regeneration and Vascular Repair and Their Application to Regenerative Medicine. Am J Pathol 191, 52–65 (2021).

49. A. R. Demonbreun et al., An actin-dependent annexin complex mediates plasma membrane repair in muscle. J Cell Biol 213, 705–718 (2016).

50. D. Bansal et al., Defective membrane repair in dysferlin-deficient muscular dystrophy. Nature 423, 168–172 (2003).

51. J. I. Greenberg et al., A role for VEGF as a negative regulator of pericyte function and vessel maturation. Nature 456, 809–813 (2008).

52. A. H. Chang et al., DACH1 stimulates shear stress-guided endothelial cell migration and coronary artery growth through the CXCL12-CXCR4 signaling axis. Genes Dev 31, 1308–1324 (2017).

53. R. S. Udan, T. J. Vadakkan, M. E. Dickinson, Dynamic responses of endothelial cells to changes in blood flow during vascular remodeling of the mouse yolk sac. Development 140, 4041–4050 (2013).

54. H. Park et al., Defective Flow-Migration Coupling Causes Arteriovenous Malformations in Hereditary Hemorrhagic Telangiectasia. Circulation 144, 805–822 (2021).

55. J. Mohammed et al., Stromal cells control the epithelial residence of DCs and memory T cells by regulated activation of TGF-β. Nat Immunol 17, 414–421 (2016).

